# Time-resolved X-ray solution scattering unveils the sequence of events leading to human Hb heme capture by *Staphylococcus aureus* IsdB

**DOI:** 10.1101/2023.08.04.551941

**Authors:** Omar De Bei, Marialaura Marchetti, Stefano Guglielmo, Eleonora Gianquinto, Francesca Spyrakis, Barbara Campanini, Stefano Bettati, Matteo Levantino, Luca Ronda

## Abstract

Infections caused by *Staphylococcus aureus* depend on its ability to acquire nutrients. One essential nutrient is iron, which is obtained from the heme of the human host hemoglobin (Hb) through a protein machinery called Iron-regulated Surface Determinant (Isd). IsdB is the protein in charge of heme extraction from Hb, which is the first step of the chain of events leading to iron transfer to the bacterium cell interior. In order to elucidate the molecular events leading from the formation of the initial IsdB:Hb complex to heme extraction, we have performed a time-resolved X-ray solution scattering (TR-XSS) investigation combined with a rapid mixing triggering approach. We succeeded in defining the stoichiometry of IsdB:Hb binding and in describing the kinetics of the subsequent structural changes. The presented approach is potentially applicable to unveil the complex kinetic pathways generated by protein-protein interaction in different biological systems.

## Introduction

The interaction between macromolecules guides and regulates all fundamental processes in living organisms. Identifying and characterizing these interactions in detail is a major challenge in life sciences, continuously calling for novel approaches. A particularly important role is played by protein-protein interactions (PPIs) in cellular processes within a given organism, or when different organisms get in contact, as in the attack of a pathogen to a host organism, where bacterial invasion and proliferation often relies on specific PPIs. A relevant example of the latter is present in *Staphylococcus aureus* infections in human hosts ^1^. One of the bacterial first needs for a successful infection is the access to iron, whose primary supply is represented by the hemic cofactor of hemoglobin (Hb). To fulfill its nutritional requirements, *S. aureus* has evolved an hemolysin arsenal to disrupt erythrocyte membrane and an Iron-regulated surface determinant (Isd) system, composed of nine proteins (IsdA-I) to bind Hb, scavenge heme and acquire iron (Fig. 1) ^2^. Several Isd proteins are cell-wall exposed and, in particular, IsdB and IsdH are in charge of extracellular Hb interception; apparently, the two receptors play a superimposable function, but only IsdB is classified as a virulence factor ^3^, eliciting a primary interest in the detailed comprehension of its mechanism of interaction with Hb. IsdB is a flexible, modular protein composed of two immunoglobulin-like domains called NEAr iron Transporter (NEAT) separated by a linker conferring mobility to the protein. The heme extraction process proceeds in two phases: the interaction between IsdB and Hb initially occurs through the NEAT1 domain and is followed by the heme transfer from Hb to the bacterial receptor, operated by its NEAT2 ^4–8^. Interfering with this mechanism is an attracting way to starve the bacterium and impair its proliferation, and represents a novel explorable target for developing antimicrobial agents against *S. aureus* and its multidrug-resistant strains, ranked by the World Health Organization as high-priority in the search of new antibiotics ^9, 10^.

**Fig. 1.**
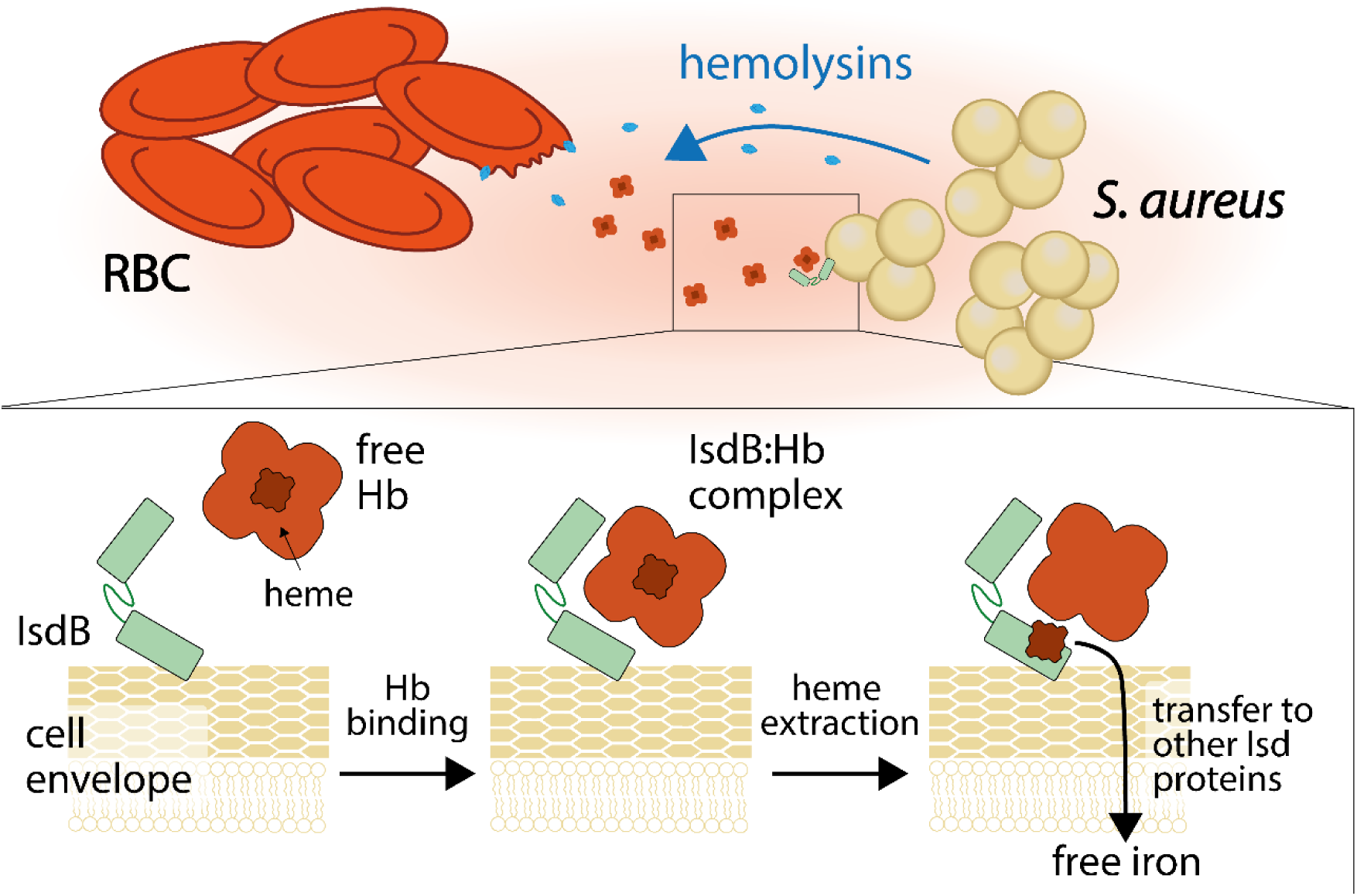
Schematic representation of the interaction between Staphylococcus aureus and human host for the bacterial acquisition of iron. The bacterium produces hemolysins to lyse red blood cells and release Hb. To proceed with the iron acquisition, S. aureus exposes the Hb receptor IsdB outside the cell wall. The magnified region shows how IsdB mediates the first step in iron acquisition: IsdB binds free Hb in the bloodstream, extracts heme and then passes it to the following proteins of the Isd system for its degradation and iron release.

The biological relevance of the PPI between IsdB and Hb motivated us in pursuing an in depth investigation of the dynamics of the whole process leading from the formation of the complex between IsdB and Hb (IsdB:Hb) to heme extraction. Up to now, this process has been extrapolated indirectly from optical spectroscopy data and available structures of IsdB:Hb complexes. By performing time-resolved X-ray solution scattering (TR-XSS) measurements after rapid mixing of *S. aureus* IsdB with human Hb we have been able to elucidate important steps in this prototypical PPI with a direct structural sensitive technique. Although XSS yields lower resolution data than X-ray crystallography, it can be applied to track in real time structural changes in solution that cannot take place within a crystal ^11^. Being directly sensitive to structural rearrangements of proteins at the tertiary and quaternary level ^11, 12^, TR-XSS has been successfully applied to investigate several biologically relevant structural relaxations including the allosteric transition of human Hb ^13–15^, the structural dynamics of light-driven proton pumps like bacteriorhodopsin or proteorhodopsin ^16, 17^, the ultrafast “protein quake” of myoglobin ^15, 18, 19^ or the photoactivation processes of phytochromes ^20, 21^ and histidine kinases ^22^. More recently, the technique has also been applied to proteins that are not intrinsically photosensitive by employing either photocaged compounds ^23–25^ or IR laser pulses to induce fast temperature jumps in the solvent ^26–29^. All of the above studies have employed a pump-probe approach with a laser pulse trigger followed by an X-ray probe pulse. An alternative possibility that could be applicable to a vastly larger class of biological systems is to follow the X-ray scattering pattern of a sample as a function of time after rapid mixing of two solutions. Although TR-XSS has been already used in combination with a stopped-flow apparatus, all studies have been so far limited to the small-angle X-ray scattering (SAXS) region ^30–34^, which mainly yields information on the overall shape of the protein and its radius of gyration. We improved the structural resolution of this approach by collecting TR-XSS data in the wide-angle X-ray scattering (WAXS) region after rapidly mixing solutions of IsdB and metHb. Indeed, this novel approach allowed us to follow, in solution and without the need of any labeling probe, the intricate sequence of protein structural changes associated with a relevant biological event. In order to extract structural information from our experimental TR-XSS signals, we have also performed TR-XSS guided molecular dynamics (MD) simulations ^35^ on the IsdB:Hb complex. We applied for the first time this approach to a protein complex, which couples the intrinsically low-resolution information from X-ray solution scattering with *a priori* physicochemical knowledge on the protein structure. The results were further validated by characterizing the interaction between Hb and IsdB through time-resolved optical absorption (TR-OA) and fluorescence (TR-F) spectroscopy in separate stopped-flow experiments. This combined approach enabled to clarify the structural organization of the IsdB:Hb complex upon the first encounter, to identify the principal kinetic steps leading to the formation of the final complex, and to estimate the corresponding kinetic rate constants. The resulting kinetic model brings into agreement the structural and functional insights separately collected so far ^6–8, 36, 37^.

Besides its relevance in biology and human health, this system is a challenging example of a PPI. In spite of the complexity of the IsdB:Hb interaction, we succeeded in obtaining a detailed description including kinetic features, stoichiometry, local conformational changes and cofactor transfer. Our combined approach, including static and time-resolved X-ray scattering and spectroscopy, together with data driven MD simulations, is applicable to a broad range of biological systems and can be used to clarify the interplay between the different processes typically occurring when biomacromolecules interact.

## Results

### Static X-ray solution scattering

We investigated the IsdB:Hb interaction both in the SAXS and WAXS regions to better characterize the sequence of events occurring when IsdB approaches Hb, and define the structure and stoichiometry of the forming complexes. SAXS allows the characterization of the particle dimensions in solution, while WAXS is sensitive to finer structural details, thus laying the basis for the interpretation of the TR-XSS data reported below. We used two different forms of Hb to isolate key intermediates along the PPI pathway, starting with formation of an IsdB:Hb complex and leading to heme extraction to the hemophore: methemoglobin (metHb, coordinating oxidized heme) and oxyhemoglobin (oxyHb, coordinating reduced heme). While metHb, which likely represents the in vivo physiological target of *S. aureus*, allows heme extraction and transfer to IsdB ^6, 8, 38^, oxyHb does not ^6^, and thus offers a snapshot of the first step of the IsdB:Hb interaction.

### Small Angle X-ray Scattering

SAXS data were collected both on IsdB:metHb and IsdB:oxyHb complexes at different concentrations (5-200 μM range). The resulting data (Fig. 2) show relatively small differences in the shape of SAXS patterns as a function of the protein concentration. While the intensity at small angles slightly increases with concentration, the measured patterns are stable in time with no indication of large aggregates in solution. Indeed, a Guinier analysis of the data shows only a relatively small increase of the radius of gyration with sample concentration (Fig S1), compatible with the presence in solution of an equilibrium between species having slightly different sizes, with larger particles more abundant at higher concentrations.

**Fig. 2.**
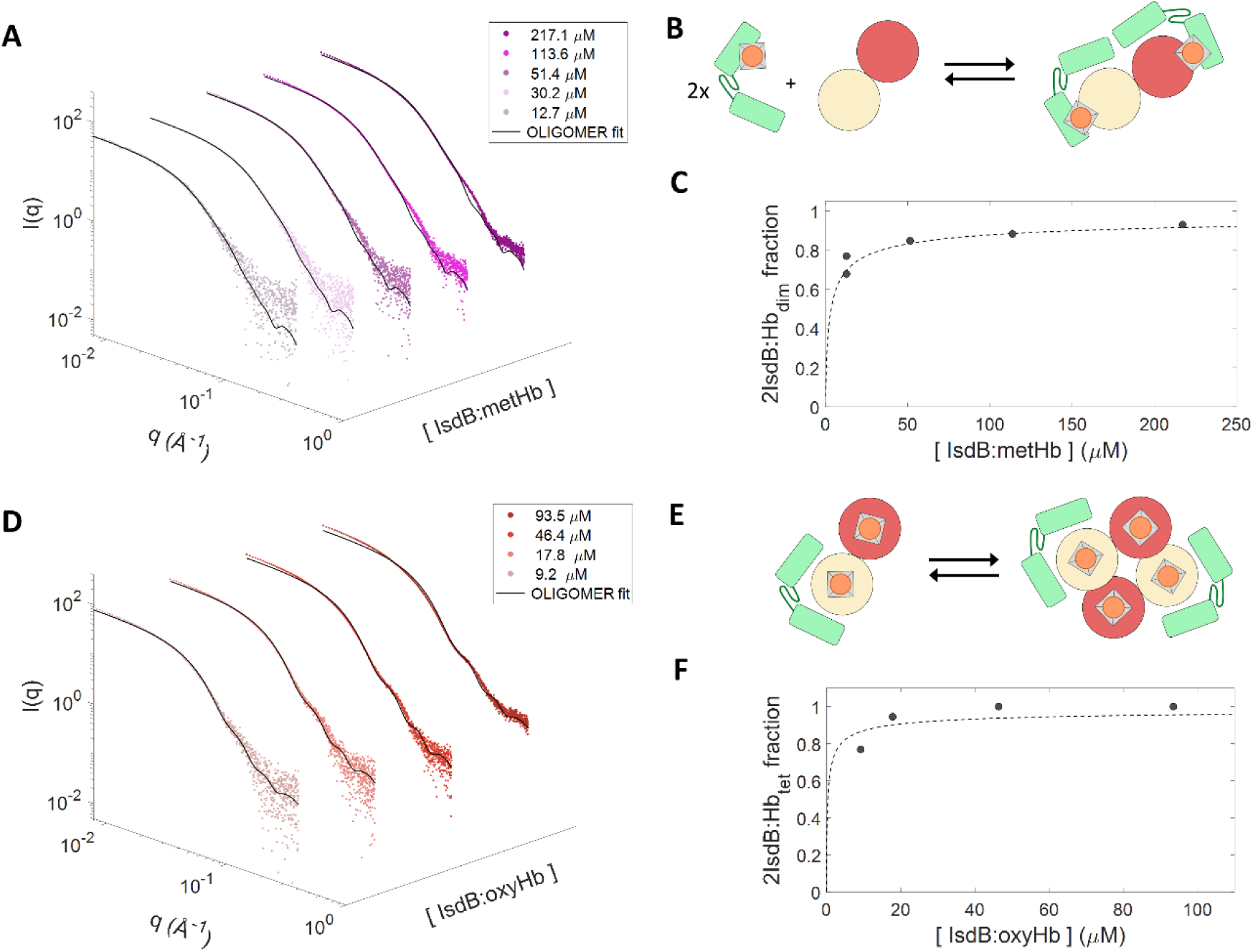
SAXS patterns of IsdB:metHb and IsdB:oxyHb complexes measured at different protein concentrations. (**A**) IsdB:metHb data (colored dots) together with fitted patterns (black solid lines) obtained with OLIGOMER. Legend reports concentrations in terms of monomeric complex (i.e. one IsdB molecule bound to a single Hb chain) (**B**) Schematic representation of the IsdB:metHb complex oligomeric equilibrium between 1IsdB:αHb_mon_/1IsdB:αHb_mon_ and 2IsdB:Hb_dim_ (**C**) Concentration of the 2IsdB:metHb_dim_ extracted from the SAXS data (closed symbols) plotted as a function of total IsdB:metHb concentration (monomeric complex). The fit (dotted line) yields a K_D_ = 1.68 ± 0.62 μM (see Materials and Methods and Supplementary Methods). (**D**) IsdB:oxyHb data (colored dots) together with fitted patterns (black solid lines) obtained with OLIGOMER. Legend reports concentrations in terms of monomeric complex (i.e. one IsdB molecule bound to an Hb dimer) (**E**) Schematic representation of the IsdB:oxyHb complex oligomeric equilibrium between 1IsdB:αHb_dim_ and 2IsdB:αHb_tet_. (**F**) Concentration of the 2IsdB:αoxyHb_tet_ extracted from the SAXS data (closed symbols) plotted as a function of total IsdB:oxyHb concentration (monomeric complex). The fit (dotted line) yields a K_D_ = 0.38 ± 1.30 μM (see Materials and Methods and Supplementary Methods).

#### IsdB:metHb complex

Comparison of the experimental SAXS patterns (and WAXS patterns, see below) with those calculated from PDB structural models of possible candidate species in IsdB:metHb mixtures (Fig. S2), allows to unambiguously conclude that, upon interaction between IsdB and metHb, a complex made of two IsdB molecules and one metHb dimer (2IsdB:metHb_dim_) is formed in solution (Fig. S3). At the highest concentration investigated, the 2IsdB:metHb_dim_ complex fully accounts for the observed X-ray pattern, while at lower concentrations an apparent lower radius of gyration is obtained, indicating the presence in solution of smaller particles. We analyzed SAXS data in terms of a linear combination of species possibly originating from 2IsdB:metHb_dim_ dissociation (carried out with OLIGOMER, see Materials and Methods). Assuming a simple equilibrium between 2IsdB:metHb_dim_, IsdB and Hb dimers, it is possible to reproduce the experimental patterns at all tested protein concentrations (Fig. 2, A and B). The resulting 2IsdB:metHb_dim_ volume fraction dependence on the total complex concentration yields a dissociation constant (K_D_) of 1.68 ± 0.62 μM (Fig. 2C). Although the resolution of SAXS data does not allow to tell whether the heme is bound to Hb or to IsdB, the time requested for sample preparation is much higher than that needed to IsdB for extracting heme from Hb, hence we concluded that dissociation mainly occurs from a complex made of holo-IsdB and apo-Hb. Indeed, dissociation from apo-IsdB and holo-Hb is known to be in the nanomolar range ^7^. Such an affinity decrease induced by heme transfer might have a physiological relevance as disengaging IsdB from Hb *in vivo* is a necessary step to allow the interaction of holo-IsdB with the following hemophore in the Isd iron acquisition pathway.

#### IsdB:oxyHb complex

SAXS patterns have been also collected for IsdB:oxyHb samples, where heme extraction does not occur, in the same range of protein concentrations used for IsdB:metHb (Fig. 2, D to F). The radius of gyration of the complex in solution at all tested concentrations is higher than the IsdB:metHb one (Fig. S1D). Also in this case, comparison with X-ray patterns calculated using PDB models allowed us to conclude that the majority of IsdB:oxyHb complexes in solution are composed by two IsdB molecules bound to an oxyHb tetramer (2IsdB:oxyHb_tet_) (Fig. S4). This is in agreement with recent cryo-EM experiments indicating that two IsdB molecules bind to the α-chains of a carbonmonoxy hemoglobin (HbCO) tetramer, when the protein concentration is relatively high (8 g/L, corresponding to 53 μM on tetramer basis) ^8^. We have fitted the SAXS pattern at the highest protein concentration as a linear combination of the scattering patterns of the 2IsdB:oxyHb_tet_ complex, and two exceeding isolated IsdB molecules to take into account the stoichiometric ratio used to prepare the sample (see Materials and Methods). In view of the cryo-EM results, the 2IsdB:oxyHb_tet_ model was built so that IsdB molecules are bound to oxyHb β-chains (2IsdB:αoxyHb_tet_). The good superposition between experimental and calculated X-ray scattering patterns (Fig. 2A) supports the hypothesis that IsdB interacts with oxyHb selectively on β-subunits, as it does in the case of HbCO. Moreover, the fitting of the scattering pattern worsens when using IsdB bound to the α-rather than to the α-Hb subunits (Fig. S5B).

The scattering patterns at the lowest concentrations deviate the most from the theoretical curve of 2IsdB:αoxyHb_tet_. Since the interaction between IsdB and Hb before heme transfer is highly stable ^7^ and their dissociation will not occur in this concentration range, the observed concentration dependence is mainly due to the dimerization of the oxyHb tetramer. We thus analyzed the SAXS patterns as a linear combination of the three following species: 2IsdB:αoxyHb_tet_, a Hb dimer with one IsdB bound to α-chains (1IsdB:αoxyHb_dim_) and isolated IsdB (Fig. 2D). The 2IsdB:αoxyHb_tet_ concentration dependence (Fig. 2F) yields a dimer-tetramer dissociation constant for oxyHb within the complex of 0.38 ± 1.30 μM. Even though the uncertainty on the estimated Hb dimer-tetramer dissociation constant is high, the best fit value is comparable to those published for Hb in the absence of IsdB in similar experimental conditions ^7^. This indicates that the binding of one IsdB to oxyHb does not appreciably destabilize the tetrameric assembly. Moreover, the fact that complexes formed by an oxyHb dimer with two IsdB do not form suggests that dimerization is a necessary but not sufficient condition for a second IsdB binding to α-subunits, as observed in metHb.

#### Wide Angle X-ray Scattering

The absence of protein aggregation over the entire range of tested concentrations was ensured by SAXS experiments, thus the highest concentration was chosen to get good signal-to-noise ratio (S/N) WAXS patterns. The WAXS patterns are useful for the interpretation of the time-resolved dataset presented below (collected over a comparable range of scattering angles), and are also functional to further validate the conclusions from the SAXS analysis. Indeed, a comparison between patterns calculated from structural models and experimental data extending up to the WAXS region is a much more stringent test than a comparison in the SAXS region only.

In order to verify that PDB atomic models of IsdB:metHb and IsdB:oxyHb complexes accurately represent the structure of the complexes in solution, we have merged the SAXS and the WAXS static data measured at the same complex concentration (≈200 μM) and compared them with the solution scattering patterns calculated from PDB models. The excellent correspondence between scattering intensities measured in the overlapping region between SAXS and WAXS, allowed us to obtain merged SAXS/WAXS patterns with remarkably good S/N in a q-range extending from 0.006 Å^-1^ to 1 Å^-1^ (Fig. 3). The calculated patterns of 2IsdB:metHb_dim_ and 2IsdB:αoxyHb_tet_ are in excellent agreement with the IsdB:metHb and IsdB:oxyHb SAXS/WAXS data, thus unambiguously confirming that these species represent the only form of the complexes present in solution (Fig. 3).

**Fig. 3.**
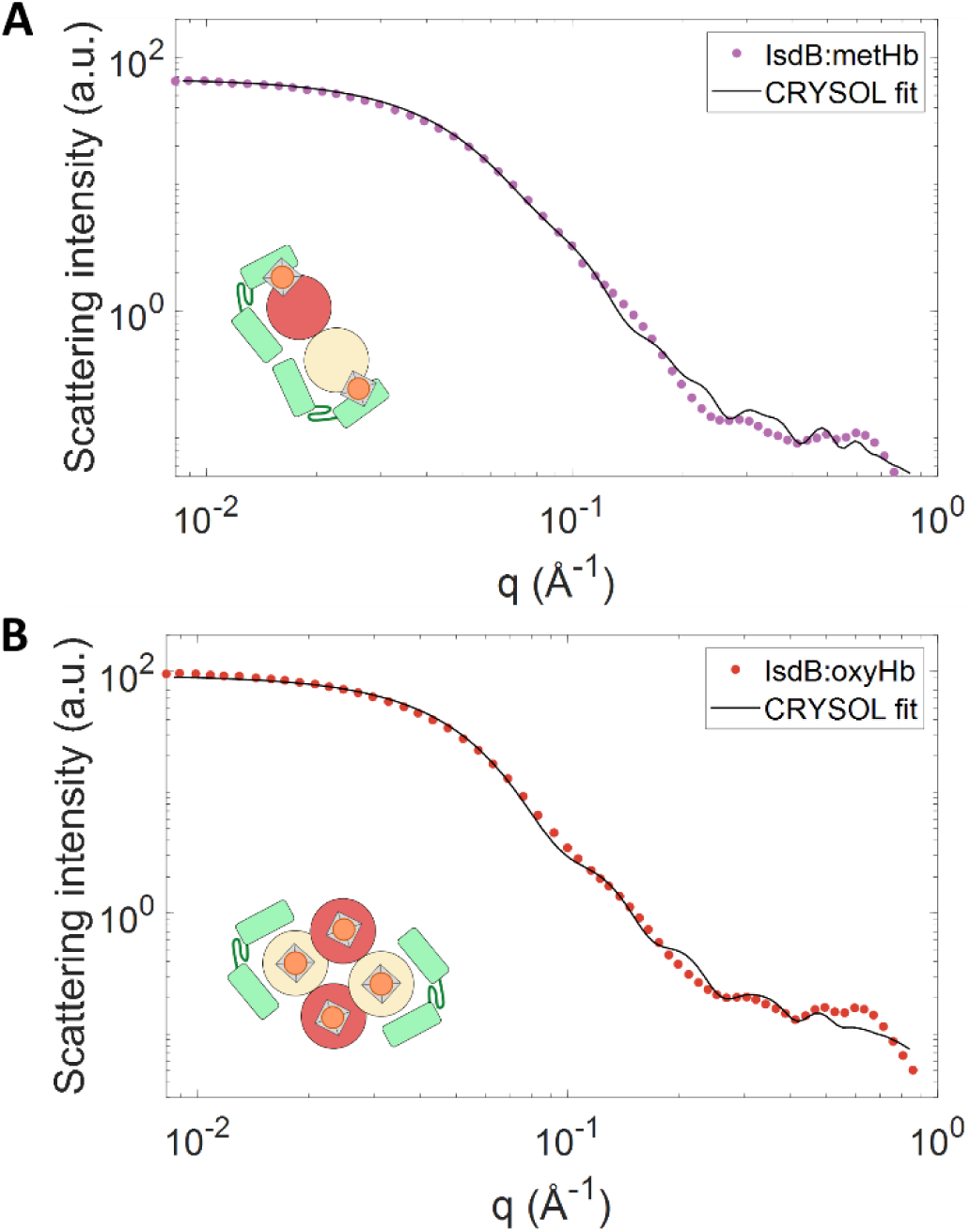
Comparison between SAXS/WAXS merged data (dots) and X-ray patterns calculated with CRYSOL. The schematic representation of the complex in solution is shown in each panel. (**A**) data relative to the equilibrium complex formed by IsdB with metHb (dotted line) vs. 2IsdB:metHb_dim_ calculated pattern (solid line). (**B**) Data relative to the equilibrium complex formed by IsdB and oxyHb (dotted line) vs. linear combination of the 2IsdB:αoxyHb_tet_ pattern and two isolated IsdB molecules (solid line, see Materials and Methods).

### Time-Resolved WAXS

The kinetics of PPI between IsdB and metHb was investigated with a new set-up for TR-XSS, employing a stopped-flow apparatus, at the ID09 beamline of ESRF (Grenoble, France) (Fig. 4). TR-XSS data have been collected in the WAXS region (TR-WAXS) after rapid mixing of IsdB and metHb solutions in equimolar concentrations (on chain basis). The total final concentration of protein was analogous to that used for static WAXS experiments (*i.e.* IsdB and metHb were mixed in an equimolar proportion; the final metHb concentration after mixing was 270 μM on a heme basis). The stopped-flow apparatus was set so that the two solutions were fully mixed and transferred to the observation capillary within 10 ms (dead time), and at each mixing event several X-ray patterns were collected at different time-delays from mixing between 10 ms and 1 min. While the observed time window does not allow to track the earlier events of IsdB:Hb interaction, it is wide enough to monitor all the structural changes triggered by the formation of the IsdB:metHb complex.

**Fig. 4.**
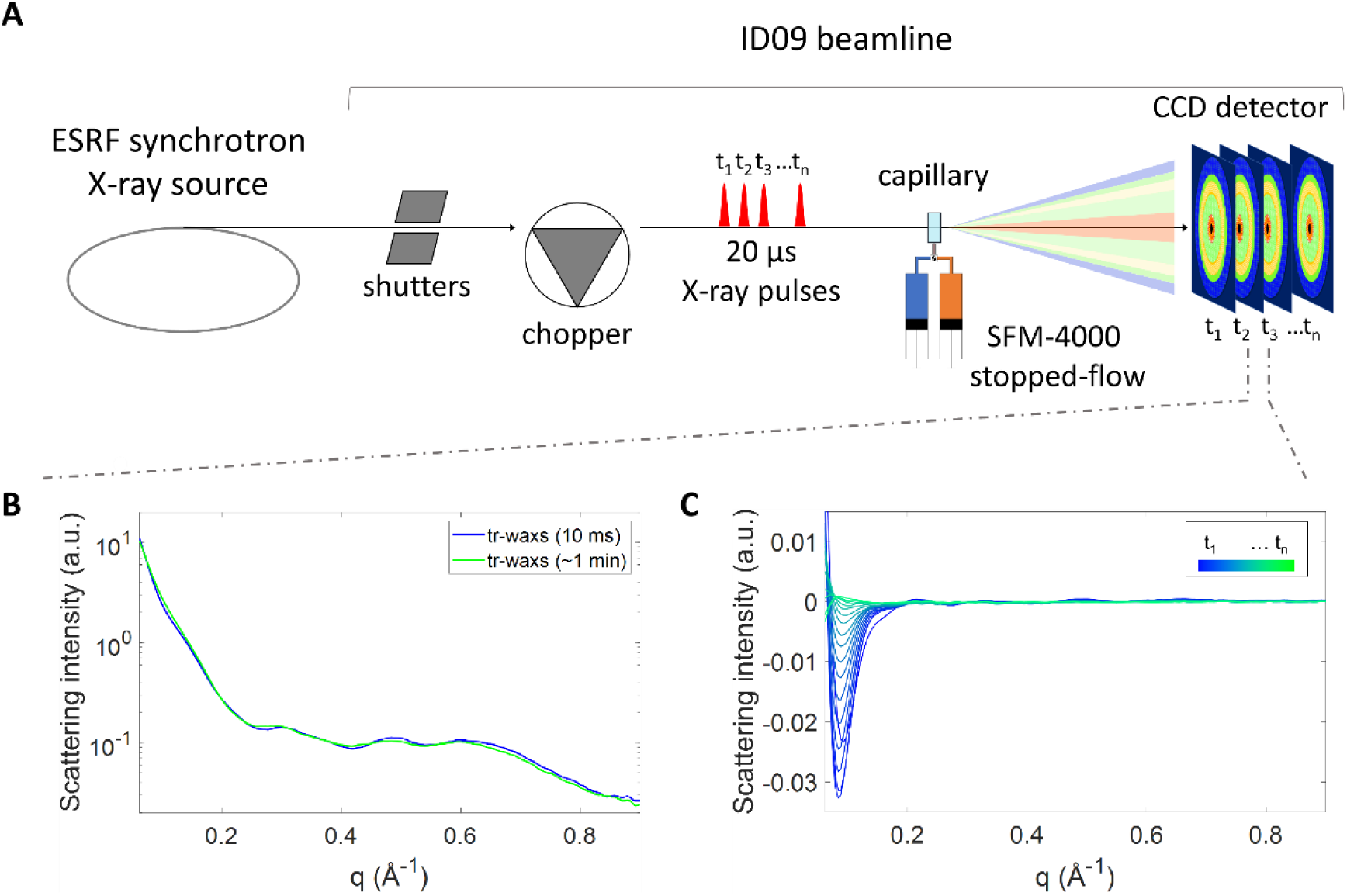
TR-XSS analysis of IsdB:metHb interaction. (**A**) Schematics of the TR-XSS setup at the ESRF ID09 beamline. X-ray solution scattering patterns are collected as a function of time after rapid mixing of an IsdB solution at a concentration of 270 μM and a Hb solution at equimolar heme concentration by means of a stopped-flow apparatus (Biologic, SFM-4000). After the two solutions were mixed and transferred to the observation capillary, induced structural changes were probed in the sample using short X-ray pulses (pulse duration = 20 μs) isolated from the X-ray source by means of shutters and choppers. The X-ray scattering pattern generated by each pulse was recorded separately through a fast CCD detector (Rayonix MX170-HS) operated in 8×8 binning in order to achieve readout times as low as 10 ms. (**B**) Comparison between the WAXS pattern measured at the earliest available time point (10 ms) and at the latest one (∼1 min). (**C**) TR-WAXS difference patterns measured at several time-delays from mixing (colormap is used to represent the time course of the reaction, where shorter delays are blue and longer are green).

As expected, the scattering patterns change within the observation time window, as shown in Fig. 4B, where first and last delays are depicted. Fig. 4C reports the time-evolution of TR-WAXS difference patterns, *i.e.* the difference between the WAXS pattern measured at time t from mixing and the WAXS pattern measured at the longest available time-delay from mixing (∼1 min). The shape of WAXS patterns changes significantly in the investigated time frame (10 ms - 1 min), with the strongest variations being around 0.08, 0.5 and 0.65 Å^-1^. We have analyzed the TR-WAXS absolute patterns (Fig. 5) in terms of a kinetic model (Fig. 5C) where the only considered processes are: (1) the sequential or concurrent binding of two IsdB molecules to a metHb tetramer (2IsdB:αmetHb_tet_), (2) the subsequent dimerization of the resulting complex in two metHb dimers each with a single bound IsdB molecule (1IsdB:αmetHb_dim_) and (3) the binding of a second IsdB molecule to IsdB:αmetHb_dim_ leading to a metHb dimer with 2 bound IsdB molecules (2IsdB:metHb_dim_). We did not consider the possibility that a complex made of a Hb tetramer with 4 bound IsdB molecules is formed as such a complex would be highly unfavorable due to steric hindrance ^6^. The symmetric binary model is based on complex stoichiometry observed in static SAXS/WAXS experiments on IsdB:metHb and IsdB:oxyHb. Indeed, according to the model, the interaction between IsdB and Hb initially leads to the formation of a Hb tetramer with 2 bound IsdB molecules, as in the equilibrium complex formed by IsdB and oxyHb (Fig. 3B), while a Hb dimer with 2 bound IsdB molecules is the final state when IsdB interacts with metHb (Fig. 3A). TR-WAXS data were fitted at each time delay as a linear combination of SAXS/WAXS patterns from static data (Fig. 5C), thus following the time evolution of the complex formation (Fig. 5B, circles). The accumulation of intermediates whose structures are different from those collected in SAXS/WAXS static experiments is expected to result in poor data fitting. Indeed, the Hb tetramer with only one IsdB bound, transiently accumulates in the early phase of the process, but cannot be isolated in solution and the corresponding static SAXS/WAXS pattern cannot be collected. For this reason, the reported fittings in Fig. 5A started from 50 ms, since after this delay this intermediate is negligibly populated (Fig. S6). Accordingly, for time delays longer than 50 ms the agreement between TR-WAXS experimental patterns and the corresponding fittings is satisfactory in the entire investigated scattering range (Fig. 5A).

**Fig. 5.**
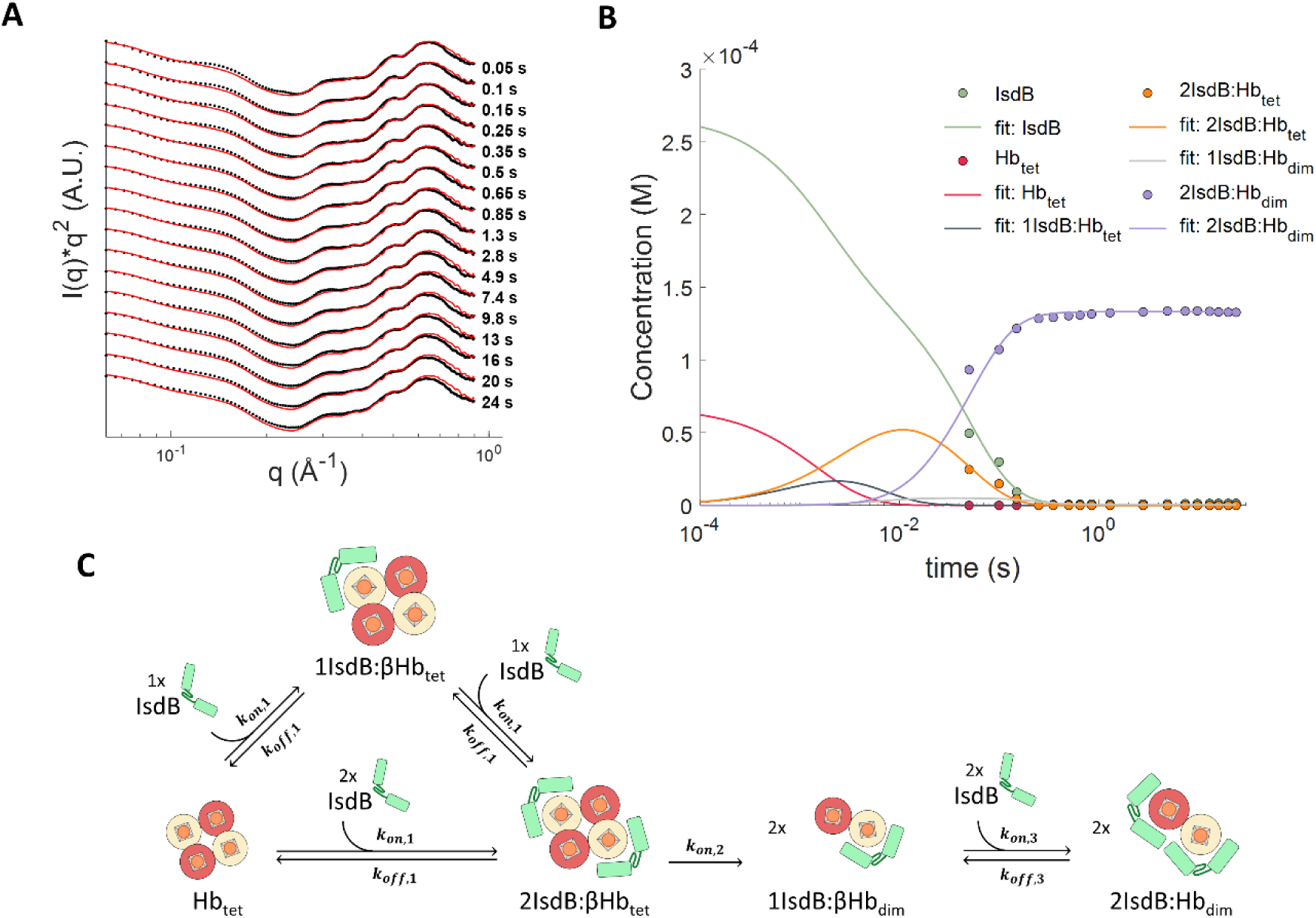
Analysis of time-resolved WAXS data in terms of a simplified kinetic model. (**A**) TR-WAXS absolute patterns (black dots) and fittings (red lines) are plotted as I(q)*q^2^ vs. q in order to amplify the differences between data and fittings at high q values. Fittings have been obtained in terms of the kinetic model described in the text and depicted in panel **C**. (**B**) Representation of species in IsdB:Hb complex formation estimated from the linear combination of TR-WAXS data (closed circles) in comparison with their global fitting (dashed lines). (**C**) Kinetic model used for the fitting the kinetic of species formation obtained from linear combination of TR-WAXS absolute patterns.

The analysis results were globally fitted to the model in Fig. 5C, allowing the determination of the microscopic constants for the described molecular processes (see Material and Methods) (Fig. 5B, dashed lines and Table 1). The calculated kinetic constants for the first IsdB binding to Hb (step 1), which well agree with published data ^7^, are also comparable to the constants for the second IsdB binding to form the final 2IsdB:metHb_dim_ complex (Fig. 5C, step 3). This evidence indicates that either Hb dimerization or activation (Fig. 5C, step 2) is the rate limiting step in the formation of the final complex. The activation process might involve a long-distance allosteric signaling that makes *⍺*-chains competent for IsdB binding with k_on_ and k_off_ values similar to those for β-chains. In fact, from the estimated kinetic parameters the resulting equilibrium dissociation constants resulted to be 7 and 3 nM for the first (to the α-chains) and second (to the *⍺*-chains) binding, respectively, in agreement with previous affinity data reported for IsdB and Hb ^7^.

**Table 1.** Kinetic constants calculated by global data fitting to the kinetic model.

|  | **STEP 1**<br>**(IsdB binding to Hbtet)** |  |  | **STEP 2**<br>**Hb**<br>**(dimerization**<br>**/ activation)** | **STEP 3**<br>**(IsdB binding to Hbdim)** |  |  |
| --- | --- | --- | --- | --- | --- | --- | --- |
| **parameter** | $k_{on,1}$ ($M^{-1} s^{-1}$) | $k_{off,1}$ ($s^{-1}$) | $K_{D,1}$ (M) | $k_{on,2}$ ($s^{-1}$) | $k_{on,3}$ ($M^{-1} s^{-1}$) | $k_{off,3}$ ($s^{-1}$) | $K_{D,3}$ (M) |
| **best fit value** | $1.4 \cdot 10^6$ | $1 \cdot 10^{-2}$ | $7 \cdot 10^{-9}$ | 20 | $2.0 \cdot 10^6$ | $0.5 \cdot 10^{-2}$ | $3 \cdot 10^{-9}$ |

### Time-Resolved Spectroscopy

Along with TR-WAXS measurements, we performed rapid mixing stopped-flow experiments collecting time-resolved optical absorption (TR-OA) and time-resolved fluorescence (TR-F) data upon mixing IsdB and metHb solutions on a benchtop instrumentation, under the same experimental conditions used to collect the TR-WAXS dataset. Visible absorbance spectra and the single-wavelength difference signal at around 400 nm (Soret peak) probe the heme cofactor and have been used to study the heme extraction process by IsdB ^5, 6^. Here, we exploited a photodiode array (PDA) detector to collect spectral evolution from 1.5 ms to the end of the heme extraction process. Singular value decomposition (SVD) was then applied to spot the two major components (Fig. S7A), which account for 80% and 11% of the signal change, and their corresponding time courses (Fig. 6A). We also followed fluorescence emission changes of tryptophans reporting variation in their microenvironment during complex formation (Fig. 6B). We preliminarily carried out static experiments on IsdB, metHb and IsdB:metHb, finding that the fluorescence signal mainly arises from IsdB tryptophans, and shows a decrease after the interaction with Hb (Fig. S7B).

**Fig. 6.**
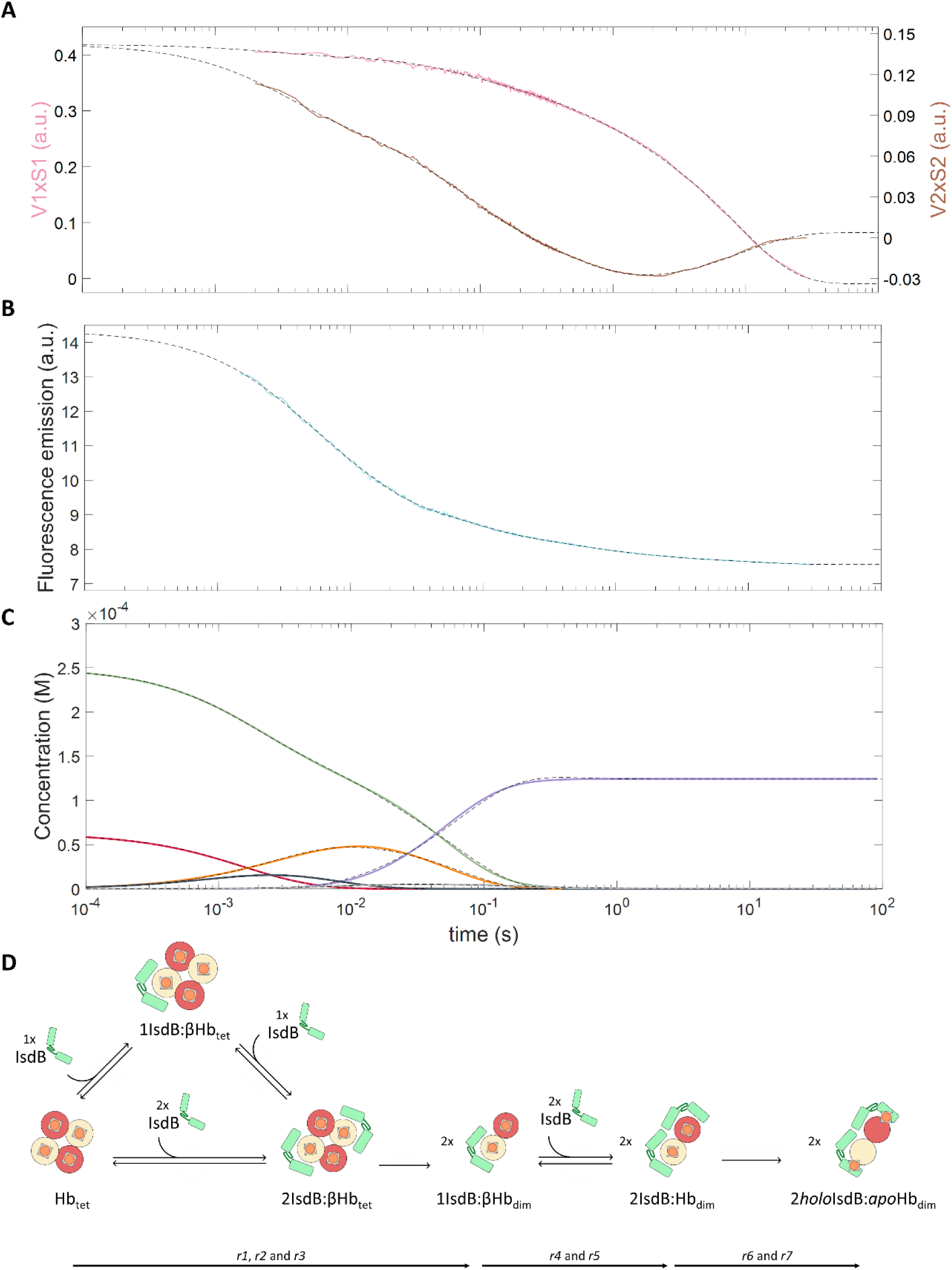
IsdB:Hb interaction followed by TR-XSS and spectroscopic techniques. Global fitting (black dashed lines) as a multiexponential function of experimental (panel **A** and **B**) and Simbiology-generated (panel **C**) data. (**A**) first (pink) and second (brown) SVD components from absorption spectroscopy. (**B**) Fluorescence (light blue) signal. (**C**) Simbiology-generated species concentrations: IsdB (green), Hb (red), 1IsdB:αHb_tet_ complex (blue), 2IsdB:αHb_tet_ complex (orange), 1IsdB:Hb_dim_ complex (grey), and 2IsdB:Hb_dim_ complex (purple). **D**. Kinetic model describing processing described by the calculated rates by global fitting analysis.

In order to extract a reliable estimation of kinetic rate constants accounting for the various processes occurring after formation of a IsdB:Hb complex (including heme extraction), we globally fit the TR-OA and TR-F kinetics together with those obtained with TR-WAXS as a sum of exponential functions sharing the same rates (Fig. 6, Table S1). Seven exponential functions were needed in order to obtain a good fit of the three datasets.

The two slowest processes (r6 and r7), that are not observed in the TR-WAXS data, account for around 80% of the first SVD component of the TR-OA data (Table S1). Since heme chromophore is the unique contributor to absorption in the analyzed spectral range, we interpret rates 6 and 7 as mainly describing heme transfer from Hb to IsdB. The fastest of the two processes (r6, 0.84 s^-1^) likely represents conformational changes in the heme environment when the final complex stoichiometry has already been reached, but heme is still bound to Hb. The slowest one (r7, 0.113 s^-1^) is in good agreement with apparent rates already attributed to the completion of the heme transfer process ^5, 7^. The faster rates (r1, r2, r3), are mainly associated with changes at the earliest time points of TR-OA data (second SVD component) and TR-F data. While the former signal could arise from IsdB binding-induced conformational changes to which the heme of Hb tetramers is sensitive, we considered IsdB tryptophan fluorescence as a marker of IsdB interaction with Hb, marginally sensitive to heme location. The remaining r4 and r5 rates largely overlap in the region corresponding to Hb dimerization/activation (step 2) and subsequent IsdB binding to *⍺*-subunits. Indeed, the fourth phase, r4, has a rate of 15.0 s^-1^ while the rate obtained from the TR-WAXS data analysis is k_on,2_ = 20 s^-1^ (Table 1).

Besides the values of estimated rates, we also considered their direction, and it is interesting to notice that the last kinetic phase in the second SVD component (r7), already attributed to heme transfer to IsdB (and kinetically-associated conformational events), shows a partial reversion, suggesting that, upon heme transfer, some conformational transitions (*i.e.* NEAT2 domain approaching and moving away from the heme pocket) partially reverse on the same conformational path.

In order to test the interpretation of the kinetic processes described above, we have performed an analogous TR-OA/TR-F stopped-flow experiment on semi-hemoglobins (semiHbs). (Fig. S8 and Table S2). SemiHbs are dimeric Hbs where heme is selectively bound only to *⍺*- (*⍺*-semiHb) or α-subunits (α-semiHb). Since semiHbs are more prone to aggregation than Hb, experiments were carried out at lower concentration (40 μM vs. 270 μM on globin basis). The time course of the first SVD component from TR-OA from both semiHbs is similar to that observed for metHb (Fig. S8 and Table S2). A large amplitude variation, attributed to heme transfer, only occurs upon IsdB binding to both α- and *⍺*-subunits, in semiHbs as well as in metHb. This evidence supports the hypothesis that IsdB likely interacts with α-subunits in all Hb variants, regardless of their holo- or apo-forms, with comparable rates, and heme extraction occurs only when both subunits in a Hb dimer are bound to IsdB.

Moreover, since semiHbs are dimeric, we demonstrated that Hb dimerization, while a necessary step for making *⍺*-subunit competent towards IsdB binding, is not the rate limiting event, but a process faster than, or rate-limited by, α-*⍺* intersubunit communication. Considering that Hb tetramer dissociation rate into dimers is in the order of 10^-3^ s^-1^ ^39, 40^, we can conclude that IsdB binding to α-subunits promotes faster metHb dimerization by several orders of magnitude, while this effect was not observed in oxyHb. These data confirmed that heme oxidation increases protein dynamics, causing a faster dimerization, and eventually activating *⍺* subunits competence for IsdB binding. Also fluorescence time courses are comparable for metHb and semiHbs, again supporting that IsdB first binds α- and then *⍺*-subunits regardless of the presence or absence of the heme cofactor (Figure S8). SemiHbs do not significantly differ from metHb in terms of rates also in the second SVD component from TR-OA, but they have a much smaller amplitude in the faster rates preceding heme transfer (Table S2). We attributed this component to dimerization or conformational changes such as interdimer activation directly sensed by the heme. In semiHbs this process is not occurring and this could explain the reduced amplitude.

### TR-WAXS driven molecular dynamics

Although the kinetic analysis of XSS and spectroscopy data allowed to clarify the sequence of kinetic events leading from the formation of a first IsdB:Hb complex to heme extraction, the analysis of TR-WAXS data as linear combination of static WAXS data does not fully exploit the structural sensitivity of the technique. To gain further insight into the details of the tertiary structural changes occurring in the observed experimental time window, we have performed molecular dynamics (MD) simulation on IsdB:Hb driven by the TR-WAXS signal. By restraining the MD simulation to conformations that are compatible with the experimental XSS patterns, it is indeed possible to obtain a more detailed structural interpretation of the experimental signals ^41^. In particular, TR-WAXS driven MD simulations have been performed against the experimental signal collected at 1.3 and 24 s from mixing. Being the system under study quite complex, the relevant settings for the simulations were chosen to avoid system instability, but to give an ensemble of configurations representing a Bayesian distribution of plausible structures (see Materials and Methods).

The TR-WAXS pattern at 1.3 s is representative of the 2IsdB:Hb_dim_ complex before heme extraction, while the TR-WAXS pattern at 24 s corresponds to the structure upon heme extraction. Unbiased MD simulations have been preliminary performed to ensure proper equilibration of the system. Then 30 ns-long TR-WAXS-driven simulations were run using GROMACS-WAXS, followed by a trajectory clustering analysis, to identify representative models of the mentioned pre-extraction and post-extraction states.

The comparison between the two TR-WAXS-driven MD simulations shows important conformational variations at different levels (Fig. 7A). The heme transfer requires the breakage of the coordination bond between the iron and the proximal histidine and, based on ^6, 8^, is also expected to be associated with the partial unfolding of the F-helix. Indeed, our simulation, driven by the TR-WAXS at the early time delay, confirms this scenario showing a general slight perturbation of both Hb chains, more evident at the α-chain, and partial unrolling of the F-helix at the C-terminus (residues 91-97 β-chain, 85-91 in *⍺*-chain). In the simulation after heme transfer, the F-helix folding is almost totally lost in α-chain, while it is preserved in *⍺*-chain. This behavior is in line with the higher dynamics of α-chain with respect to *⍺*-chain in metHb ^42^. Moreover, a roto-translation movement involving linker and NEAT2 can be observed at the level of both IsdBs promoting the detachment from Hb (Fig 7, C and E). This is not surprising, since, upon F-helix unfolding and/or coordination loosening, the heme can undergo significant rotation and translation and be more easily pulled by the NEAT2 domain.

**Fig. 7.**
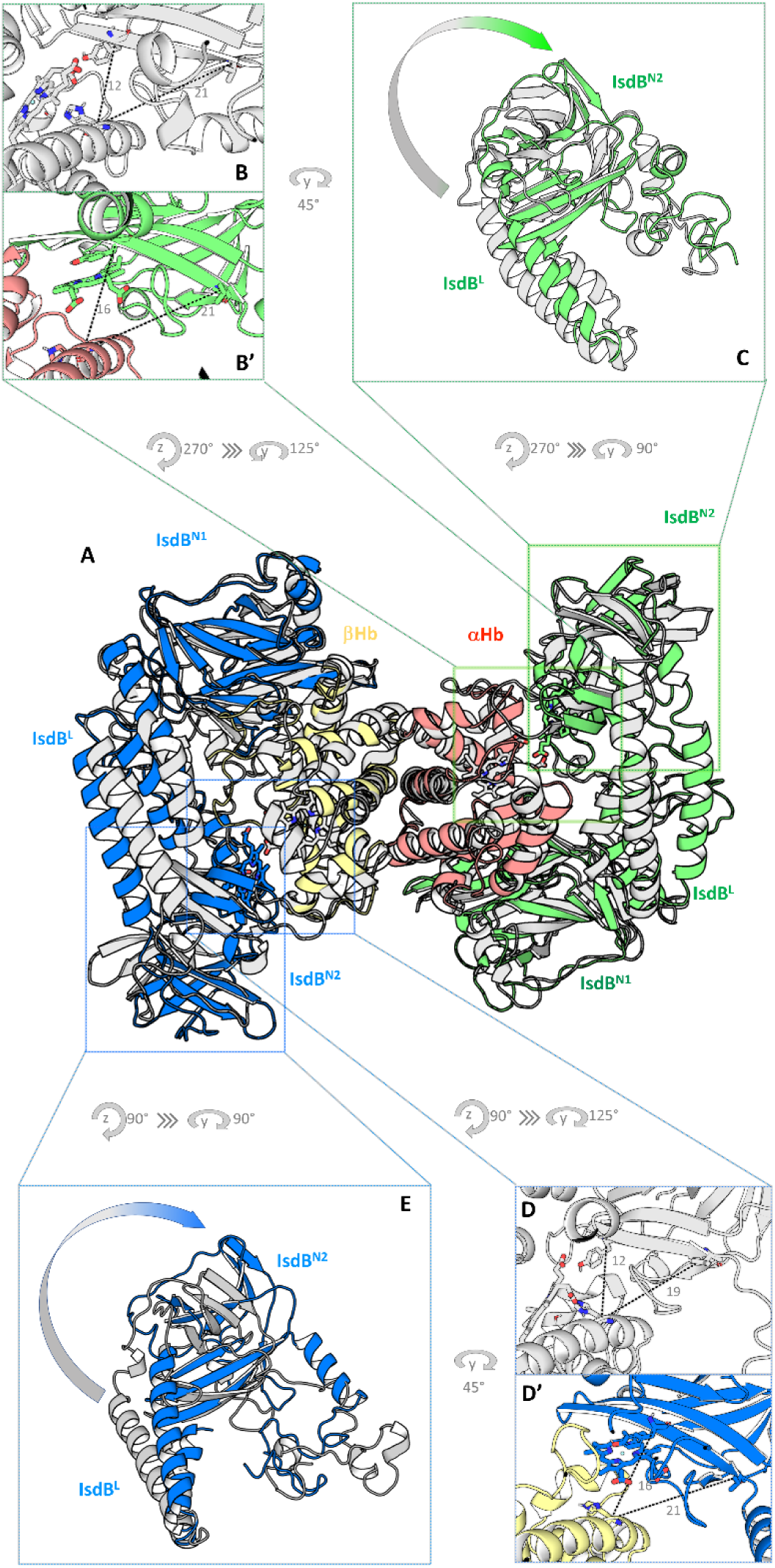
Investigating pivotal steps in IsdB:metHb interaction through XSS-driven MD simulations. (**A**) Superposition of md1.3 and md24 complexes (gray and colored cartoons, respectively). (**B**) His(C*⍺*)-Tyr444(C*⍺*) and His(C*⍺*)-Val449(C*⍺*) distances measured for Hb *⍺*-chain and IsdB NEAT2 in md1.3 complex. (**B’**) His(C*⍺*)-Tyr444(C*⍺*) and His(C*⍺*)-Val449(C*⍺*) distances measured for Hb chain a and IsdB NEAT2 in md24 complex. (**C**) Roto-translation underwent by linker and NEAT2 domain of IsdB when interacting with Hb *⍺*-chain. (**D**) His(C*⍺*)-Tyr444(C*⍺*) and His(C*⍺*)-Val449(C*⍺*) distances measured for Hb α-chain b and IsdB NEAT2 in md1.3 complex. **D’**. His(C*⍺*)-Tyr444(C*⍺*) and His(C*⍺*)-Val449(C*⍺*) distances measured for Hb α-chain and IsdB NEAT2 in md24 complex. (**E**) Roto-translation underwent by linker and NEAT2 domain of IsdB when interacting with Hb α-chain. Proteins are represented in cartoons, while heme groups are depicted as sticks. Distances are reported in Å.

Domain movements between the two snapshots obtained from TR-WAXS driven MD simulations were compared by measuring distances between representative residues. In particular, to reveal the effect of the roto-translation, the distance between Hb proximal histidines (His87 on β-chains, His92 in *⍺*-chains) and two different IsdB residues has been measured. In β-chain, the distance between the alpha carbons of Hb His87 and IsdB Tyr444 changes remarkably by 4 Å (from 12 to 16 Å), while the distance between Hb His87 and IsdB Val449 change by 2 Å (from 19 to 21 Å) (Fig. 7, D and D’). In the case of *⍺*-chain, the variation is detected only at the level of Hb His92 and IsdB Tyr444 distance, increasing from 12 to 16 Å (Fig. 7, B and B’). Despite the different Hb chain dynamics, the IsdB roto-translation appears to follow the same pathway, thus suggesting a similar mechanism of detachment of linker and NEAT2 domains from Hb.

The analysis of TR-WAXS driven MD trajectories allows the identification of transient conformations that were not previously observed. In particular, it was possible to obtain information on the structural changes preceding heme extraction and leading to the affinity decrease between the two proteins.

## Discussion

By integrating static and TR-XSS, TR absorption and fluorescence spectroscopy, and MD, we were able to dissect the complex stoichiometry, the Hb oligomerization state, and the kinetics along the reaction pathway following the interaction between IsdB and Hb and leading to heme extraction: IsdB binding to oxyHb β-subunits does not significantly destabilize Hb tetramers, and dimerization is a necessary but not sufficient condition for a second IsdB binding. This latter event only occurs on metHb, where intersubunit communications triggered by IsdB binding to α-chains make α-subunits competent for IsdB binding. Only when both globin chains on a Hb dimer are bound to IsdB, heme extraction proceeds and the cofactor transfer is coupled to linker-NEAT2 domain movement. F-helix refolding in α-chains, but not β-chains, supports a different dynamics of the two subunits.

We challenged our kinetic model and microscopic constants for their ability to predict how IsdB and Hb interact in different experimental conditions. The good agreement (Fig. S9 and Table S3) between the rate constants from our simulation and published ones ^6^ proves the approach we developed can be confidently used to describe *in vitro* behavior of IsdB:metHb PPI. These findings can be evaluated in view of their physiological meaning and implications. Hb free in the plasma from hemolyzed red blood cells circulates mainly as dimers, and IsdB can bind dimers or tetramers on β-subunits with the same efficiency. Binding of a second IsdB molecule present in a spatial proximity is then required to trigger concomitant heme extraction. This would guarantee a fully productive interaction of the bacterium with the iron containing protein. This mechanism implies that IsdB molecules are anchored to the cell wall at a distance suitable for forming a ternary complex. This could be expected considering the necessary colocalization of other proteins of the Isd system, such as IsdA, to allow heme transfer and internalization, and the observed surface distribution of IsdB (and IsdA) under iron starvation conditions that localize to discrete puncta throughout the cell surface ^43^. Besides biological implications of this mechanism, these findings possibly contribute to the design of inhibitors of the interaction between IsdB and Hb as a way to bacterial iron starving: based on the identified mechanism, molecules binding on one of the two globin chains would likely affect heme extraction.

To our knowledge, this is the first study reporting coupling of rapid mixing TR-WAXS with structural and spectroscopic analyses, which can attribute structural determinants to microscopic kinetic constants, applied not to a single molecular entity but to a higher complexity system, *i.e.* the characterization of structural dynamics following protein-protein interactions.

## Materials and Methods

### Proteins expression and purification

Expression and purification of StrepTag®II-IsdB and purification of human Hb were carried out as previously described ^7, 8^. All protein samples used for the analysis were dialyzed against the experimental buffer containing 100 mM Tris/HCl, 150 mM NaCl, and 1 mM EDTA, pH 8, flash-frozen in liquid nitrogen, and stored at -80 °C. SemiHbs were prepared as published elsewhere ^7, 44^ and detailed in Supplementary materials.

### Static X-ray solution scattering data collection

Static SAXS and WAXS data were collected at the BM29 beamline and at the ID09 beamline of the ESRF, respectively. Several preliminary measurements were carried out to find optimal working conditions in terms of solvent composition and pH. The exposure time has been adjusted to make sure that X-ray-induced radiation damage was negligible. For SAXS data collection an X-ray energy of 12.5 keV and a sample-to-detector distance of 2.849 m were used, while WAXS data were collected at 17.45 keV and 0.39 m. In both cases, approximately 50 μl of each sample was loaded into a 1.7 mm quartz glass capillary. Immediately before the data collection, samples were centrifuged at 17,200 g for 30 min at 4 °C and the supernatant was collected and loaded into the sample exchanger after checking its optical absorption spectrum with a Nanodrop spectrometer (Thermo Scientific). The protein concentration ranged from 12 to 220 μM for SAXS and from 90 to 270 μM for WAXS measurements. SAXS data were collected using a Pilatus 1M detector at an exposure of 0.5 s per image while the sample was flowing through the capillary at a flow speed of 5 μL/s. WAXS data were collected using a Rayonix MX170-HS detector exposing the sample to short (∼3 μs) X-ray pulses at a repetition rate of 10 Hz while the sample was translated in steps of 60 μm in between two consecutive X-ray pulses. Microsecond X-ray pulses were selected using a high-speed chopper operated in the so-called tunnel-less mode ^45^. Typically, 10 images per sample were azimuthally integrated using pyFAI in order to obtain the scattering intensity as a function of the scattering vector magnitude q=4πsin(θ)/λ, where 2θ is the scattering angle and λ is the X-ray scattering wavelength. Scattering patterns from the same sample were averaged together after verifying that they were perfectly superimposable. Buffer data were collected before and after each protein sample and used as a reference to calculate the protein excess scattering signal. The protein radius of gyration (R_g_) was estimated from the SAXS data either by the automatic software available at the BM29 beamline or by custom made python scripts. WAXS data were collected at the highest protein concentration at which SAXS data show minimal sign of protein aggregation or interparticle interference effect. SAXS and WAXS data were then merged and rebinned so that experimental points were homogeneously spaced on a logarithmic q-scale.

### Modeling of X-ray solution scattering data

In order to extract structural information from the SAXS/WAXS data we compared experimental patterns with curves calculated from structural models using CRYSOL 3.0 ^46^. We considered all possible *a priori* candidate models (Fig. S2) of the IsdB:Hb complex starting from a metHb monomer bound to a single IsdB molecule up to a Hb tetramer bound to four IsdB molecules. All models have been generated starting from the available crystallographic structure of *S. aureus* IsdB bound to human Hb (PDB ID 5VMM) where the Hb component was replaced by a high resolution metHb crystallographic structure (PDB ID 3P5Q). The I-TASSER web server ^47^ was used to model the missing StrepTag®II in the 5VMM PDB model. A full metHb tetramer with 4 bound IsdB molecules (4IsdB:Hb_tet_) was thus generated and all subsequent models were obtained by removing elements. Thus, the 2IsdB:Hb_dim_ was generated by removing one Hb dimer with the two bound IsdBs, the 1IsdB:Hb_dim_ was generated by removing an additional IsdB and so on. By directly comparing the experimental SAXS/WAXS patterns with each of the above models (Fig. S3 and S4), it was possible to exclude several possible candidate structures.

### Fitting of SAXS dilution series

An overall good agreement with the IsdB:metHb SAXS dilution series (Fig. 2A) can be obtained using a model containing a metHb dimer and 2 bound IsdB molecules (Fig. S2). However, small deviations were observed especially when the calculated patterns were compared with the data measured at the lowest protein concentration. We have fitted the SAXS data, to consider possible dissociation equilibria, as a linear combination of IsdB, Hb_dim_, 1IsdB:*⍺*Hb_mon_, 1IsdB:αHb_mon_, IsdB:Hb_dim_ models (Fig. S2) using OLIGOMER ^48^. Volume fractions of models were constrained so that 2IsdB:metHb_dim_ dissociate into one Hb dimer and two IsdB molecules or IsdB bound to single Hb chains.

Data corresponding to IsdB:oxyHb sample were found to best fit using models made of two IsdB molecules bound to the *⍺*- or the α-chains of an Hb tetramer (2IsdB:αHb_tet_ or 2IsdB:αHb_tet_), with model 2IsdB:αHb_tet_ giving a best far agreement (Fig. S5B). Even in the SAXS series of this complex data measured at lower protein concentration showed deviations from the theoretical curve. In this case, OLIGOMER fits of the SAXS data at different concentrations have been performed as linear combinations of the above pattern (model 2IsdB:αHb_tet_), the pattern of one IsdB bound to the α-chains of an Hb dimer (model 1IsdB:αHb_dim_), and the pattern of isolated IsdB. The reason why this latter structure was included is to consider the 1:1 globin chain:IsdB stoichiometric ratio of the mixture prepared for scattering experiments, and the same ratio was constrained throughout OLIGOMER analysis (namely one isolated IsdB for each 1IsdB:αHb_dim_ and two isolated IsdB for each 2IsdB:αHb_tet_). Also in this case, adding models of the isolated proteins or the complex in a different stoichiometric ratio (namely model 2IsdB:Hb_dim_) (Fig. S5A) did not improve OLIGOMER fitting.

Finally, theoretical patterns of the above selected models or model mixtures were compared to the merged SAXS/WAXS patterns of both IsdB:metHb and IsdB:oxyHb, which define the analyzed sample with higher resolution, to further confirm the achieved results. SAXS concentration series of both IsdB:Hb complexes were fitted using OLIGOMER to determine the change in the oligomeric state of Hb within the two complexes (IsdB:metHb and IsdB:oxyHb). The dissociation constant for the above equilibrium was calculated adapting the equation previously reported previously ^49^, see Supplementary Methods.

### TR-WAXS data collection

TR-WAXS data were collected at the ID09 beamline of the ESRF. Samples were prepared similarly to those used for the static SAXS and WAXS data collections. Solutions of human metHb and *S. aureus* IsdB at equimolar concentration of 267 μM were loaded into two different syringes of an SFM-4000 Bio-Logic stopped-flow apparatus equipped with a 1.4 mm quartz glass capillary. In order to get a final equimolar concentration, 28 μL of the metHb solution was mixed with 72 μL of the IsdB solution. The flow rate of mixing (1.67 mL/s) was optimized in order to minimize any inhomogeneity of the sample and corresponds to a dead time of 10 ms. X-ray scattering patterns in the 0.05-2.2 Å^-1^ scattering region were recorded with a Rayonix MX170-HS detector at an X-ray energy of 17.67 keV and a sample-to-detector distance of 0.4 m. Each image is the result of the interaction of the sample with a single 20 μs X-ray pulse. At each mixing event, up to 24 images were recorded at different time delays from mixing between 10 ms and 1 min. By operating the Rayonix detector in frame-transfer mode and 8×8 binning, it was possible to record consecutive images every 30 ms in the fast part of the kinetics. Preliminary measurements were used to optimize the exposure time per image and the total number of images per mixing event so as to minimize any heating or radiation damage effect on the sample. TR-WAXS images were azimuthally averaged using pyFAI and the resulting 1D curves were normalized in the 2.1-2.2 Å^-1^ water scattering region before further processing to correct for fluctuations in the intensity of the X-ray beam ^13^. The data collection sequence was repeated for at least 50 times and TR-WAXS patterns at the same time-delay were averaged together to increase the signal-to-noise ratio of the dataset.

The resulting S(q, t) patterns were used to calculate TR-WAXS difference patterns as ΔS(q, t) = S(q, t) - S(q, 1 min), *i.e.* the WAXS pattern measured at the longest available time delay (1 min) was used as a reference. TR-WAXS absolute patterns were instead obtained as S(q, t) - S_buffer_(q), where S_buffer_(q) is the equilibrium WAXS pattern of the buffer solution measured with the same stopped-flow apparatus used for the TR-WAXS data collection. TR-WAXS absolute patterns are thus proportional to the excess protein scattering contribution and directly comparable to the static XSS data.

### TR-WAXS data analysis

The data analysis reported in the main text is focused on the 0.01-1 Å^-1^ q-range where pattern changes are expected to originate mainly from stoichiometry changes and tertiary rearrangements. Patterns changes in the 1-2.2 Å^-1^ range are dominated by changes in the solvent structure around the IsdB:Hb complex induced by heat transfer between the protein complex and the solvent (Fig. S10).

The theoretical WAXS pattern for each time delays, S_theo_, can be written as linear combination of experimental static WAXS scattering patterns, S, of isolated proteins, IsdB:oxyHb complex, and IsdB:metHb complex:

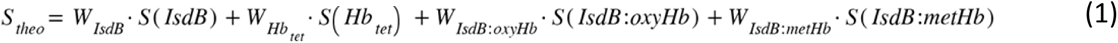

Weights vectors for each molecular species over the time were obtained using a least-squares fit by comparing experimental TR-WAXS pattern for each time delay and theoretical pattern as described above, S_theo_, using Fminuit minimization algorithm in MATLAB. The minimization was only constrained to keep weights of isolated proteins constant over time to take into account the stoichiometric ratio used to prepare the sample. During the fitting step, weight of IsdB was calculated as follows:

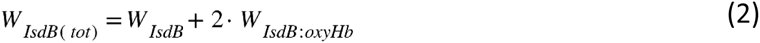

because the scattering pattern of IsdB:oxyHb complex describes a sample where a mixture of 2IsdB:Hb_tet_ complex and 2-times molar excess of IsdB are present in solution (as discussed in the main text). Weights were normalized by dividing each value by the one of IsdB:metHb complex at the last delay (which is the only species present at the end of the reaction). Normalized weights were then multiplied by 135 μM, which is the concentration of the final complex in our experimental conditions, to obtain concentrations of different species over the time. This linear combination was carried out on the complete dataset and the resulting weights vectors over the time were used as input to SimBiology ^50^ to extrapolate microscopic constants for the molecular processes involved in IsdB:metHb protein-protein interaction. We performed the global fit of TR-WAXS dataset employing non-linear regressions statistical model, Fmincon as local solver and scatter search as global solver, a proportional error model for each molecular species and by pooling all estimated parameters together. SimBiology was also used to simulate the concentration of molecular species over time.

### TR-OA and TR-F spectroscopy

TR-OA and TR-F experiments were carried out using an SX18 stopped-flow apparatus (Applied Photophysics) with a dead time of about 1.5 ms equipped with a 150 W xenon lamp and alternatively coupled to a photomultiplier for fluorescence spectroscopy or to an SX Direct Coupled Photodiode Array Detector for multi-wavelength absorption measurement. The kinetics of the reaction between equimolar amounts of IsdB and metHb (globin concentration) at concentration mimicking TR-WAXS experiments (250 μM of IsdB and metHb) or at lower concentration (40 μM IsdB and Hb) were measured at 20 °C in the experimental buffer. The interaction between IsdB and semiHbs was assessed to kinetically evaluate whether the hemophore interaction with Hb is affected by the absence of heme. Since semiHbs appeared unstable under the TR-WAXS experimental conditions (namely, 250 μM globin concentration), we tested this reaction by mixing equimolar concentrations of IsdB and semi(*⍺*) or semi(α)metHb at a concentration of 40 μM.

Fluorescence emission (298 nm excitation) was recorded at 90° after passing through a 320 nm cut-off filter by acquiring 10,000 data points for up to 30 s. For rapid-scanning experiments, 1,000 logarithmically spaced spectra of the reaction mixture (in the 320–700 nm interval) were collected for up to 30 s. Since the concentration used for TR-WAXS experiments was too high to follow the reaction over the entire range of selected wavelengths, we focused our analysis on Q bands (525– 700 nm interval) that have suitable molar extinction coefficients.

### Global fitting of kinetic data to a multiexponential model

Global fitting was performed adapting lsqmultinonlin function ^51^. Time courses of concentration for molecular species simulated by SimBiology Model Analyzer, absorbance and fluorescence spectroscopy were fitted with a sum of seven or six exponential functions. The amplitudes of the signals were left free to change while the exponential rates (r1 to r7) were kept the same throughout the entire dataset.

### TR-WAXS driven MD

Two different TR-WAXS driven MD simulations have been performed against either the experimental pattern measured at 1.3 s from mixing or that at 24 s from mixing. The pattern at 1.3 s is representative of the 2IsdB:Hb_dim_ complex before heme extraction and the structure used for the simulation is exactly the one described in the main text and used for the SAXS data analysis. The pattern at 24 s is representative of the structure after heme extraction has occurred, and for the simulation we used the low-resolution structure solved by cryo-EM (PDB ID 7PCF, ^8^), to which the StrepTag®II sequence at the C-terminus was added with the I-TASSER web server ^47^. As a preliminary preparatory step, an unbiased MD simulation was carried out using GROMACS 2022.1^52^ starting from either of the two above models. The systems were parameterized using amber99sb-ildn force field ^53^ for protein; water and ions were treated using the TIP3P model ^54^. Proteins were soaked in a dodecahedral water box, with a 30 Å pad and neutralized with NaCl. Short range interaction cut-off was set at 10 Å. Long-range electrostatic interactions were treated with Particle Mesh Ewald algorithm ^55^. Temperature control was achieved using the modified Berendsen thermostat ^56^. Minimization and equilibration was achieved through different steps: 2.5 ns equilibration in the NVT ensemble at 100 K with heavy atoms restrained; 2.5 ns equilibration in the NVT ensemble at 100 K without restraints; 2.5 ns equilibration in the NVT ensemble at 300 K; 2.5 ns equilibration in the NPT ensemble at 300 K with heavy atoms restrained and 2.5 ns equilibration in the NPT ensemble at 300 K without restraints using Berendsen barostat ^57^ and finally 2.5 ns equilibration in the NPT ensemble at 300 K using Parrinello-Rahman barostat ^58^. In all steps the integration time step was set to 1 fs^-1^. The systems were then subjected to 20 ns of plain MD run, using a time step of 2 fs^-1^, and the obtained configurations were used as starting points for XSS-driven MD.

The biased simulations were carried out using a modified version of GROMACS 2016 (GROMACS-SWAXS). The systems were simulated in the NPT ensemble for 30 ns at 300 K, with a time step of 2 fs^-1^. For the biased simulation, the relevant settings were chosen as follows: time constant *waxs-tau* for averaging on-the-fly calculated curves was set to 500 ps; *waxs-t-target* was set to 5000 ps, to gradually turn on the waxs derived potential; waxs potential was linearly coupled to experimental scattering data and the force constant was set to 1: in such a framework the ensemble generated by WAXS-driven MD simulation can be interpreted as the posterior distribution of a Bayesian inference problem, given experimental data (WAXS pattern) and prior physical knowledge (the force field used for simulation) ^59, 60^; with the same aim, *waxs-solvdens-uncert-bayesian* was set to “yes”, in order to sample error of the solvent density in a Bayesian manner, simultaneously with the refined structure.

The portion of the trajectories displaying a stable biasing potential was used for the subsequent analyses. In general, the conformations sampled during simulations are not significantly different with respect to the starting models, thus supporting the reliability of the applied methodology. Relevant configurations were extracted through gromos clustering algorithm ^61^, as implemented in GROMACS package. The different clusters were analyzed by identifying their medoids (cluster centroids) and classified on the basis of their conformation population. Among the most populated clusters, the medoids have been further scrutinized on the basis of a priori psycho-chemical knowledge on the IsdB:Hb complex. In particular, we have selected the most populated cluster with the additional requirement that the distance between Hb and IsdB was compatible with bond formation. Since it is well known that the IsdB NEAT1 domain is responsible for the majority of the hemophore affinity for Hb ^4^, this distance requirement was applied only to the IsdB NEAT1 domain.

## Acknowledgments

The X-rays solution scattering experiments have been performed at the European Synchrotron Radiation Facility (ESRF): static and time-resolved WAXS experiments were performed at the ID09 beamline (experiments IH-LS3074 and LS-2807), while the static SAXS experiments were performed at the BM29 beamline. Petra Pernot and Gabriele Giachin are kindly acknowledged for assistance in using beamline BM29. The Partnership for Soft Condensed Matter (PSCM) is also kindly acknowledged for support during the experiments at the ESRF. We thank Dr. Manuele Bettelli (IMEM-CNR) and Prof. Stefania Abbruzzetti (University of Parma) for helpful discussion and technical support preparing MATLAB scripts.

