## Supplementary materials for "Time-resolved X-ray solution scattering unveils the sequence of events leading to human Hb heme capture by *Staphylococcus aureus* IsdB"

### Supplementary Text

#### IsdB expression and purification

The sequence of the gene coding for the central and functional region of IsdB (UniProt entry Q8NX66), residues 125-485, was optimized for expression in *E. coli* and subcloned in pASK-IBA3plus (IBA Lifesciences) vector with a C-terminal StrepTag®II. Protein overexpression was carried out in *E. coli* cells in LB broth medium at 37 °C, expression was induced with 0.2 µg/mL anhydrotetracycline at the mid-log phase ($OD_{600}$ = 0.5 – 0.6) followed by incubation at 20 °C for 24 h. The purification of the overexpressed protein was performed on a pre-equilibrated column, packed with Strep-Tactin® XT resin (IBA Lifesciences), and a final SEC step (HiLoad 16/600 Superdex 75 prep grade column, GE Healthcare) was performed to remove high MW contaminants. The final protein preparation was over 95% pure, and the amount of holo-IsdB, estimated using a published extinction coefficient of 90,500 $M^{-1}$ $cm^{-1}$ at 405 nm [7], was lower than 5% of the total amount of the protein.

#### Hb purification

Human Hb A was purified from outdated blood from non-smoking donors obtained from a local blood transfusion center. The supernatant containing oxyHb was separated from cell debris of lysed red blood cells (RBC) by centrifugation, for 1 hour at 4 °C and 23,000 g. Hb was purified from soluble RBC components using a 100 x 5 cm CM-Sephadex C-50 column, following a linear pH gradient from 6.8 to 8.6. Visible absorption spectroscopy was used to calculate the concentration and to evaluate the oxidation state of purified oxyHb [62].

Oxidation of oxyHb to metHb was achieved by using potassium ferricyanide (Fluka). Hb was incubated for 5 minutes in the presence of the oxidizing agent at 5 mM concentration, then the latter was removed on a G-25 Sephadex desalting column (GE Healthcare). The metHb concentration was calculated spectroscopically by diluting the protein in the experimental buffer (100 mM Tris/HCl, 150 mM NaCl, and 1 mM EDTA, pH 8), where the extinction coefficient at 406 nm is known ($\varepsilon_{406}$ = 130,000 $M^{-1}$ $cm^{-1}$) [8].

#### SemiHbs preparation

SemiHbs were prepared by mixing apo-Hb with isolated α- and β-subunits. The reaction was completed in 65 h at 4 °C and the desired product was separated from unreacted species with a final chromatographic step using a GE Healthcare HiLoad 16/600 Superdex 75 prep grade column. To prepare apo-Hb, metHb is added to an equal volume of butanone in a separatory funnel at 4 °C and the pH was adjusted to 2.5 with 1 M HCl under vigorous stirring. After standing 1 min at 4 °C, the aqueous phase was separated from the organic layer. Acidic solution was intended to denature Hb resulting in the heme loss, with the latter moving into the organic solvent. The resulting solution was exhaustively dialyzed against distilled water to remove butanone, then against 0.2 M phosphate buffer pH 7.0, and finally against the buffer used for semiHbs preparation (0.1 M phosphate pH 7, 150 mM NaCl, 1 mM EDTA). The preparation of separated α and β subunits in their native state was carried out by following the method from Bucci and Fronticelli [63]. Briefly, HbCO was mixed with p-chloromercuribenzoate (CMB) in 20 mM potassium phosphate pH 7 at 1:8 Hb:CMB ratio. Ionic strength was brought to 0.1 M with potassium chloride and the pH was adjusted to pH 6, then reaction was completed overnight at 4 °C. The resulting solution was equilibrated in 10 mM potassium phosphate pH 5.8 and loaded on a CM-Sephadex column. The elution of separated Hb subunits was achieved with a pH gradient, β-Hb chains eluted at pH 6.4 while α-Hb chains eluted at pH higher than 7.4. After the separation, CMB was removed from Hb subunits by adding cysteine at 50-molar excess with respect to the number of thiol groups to be regenerated. Finally, protein solutions were exchanged in the buffer needed for semiHbs preparation (0.1 M potassium phosphate pH 7, 150 mM NaCl, 1 mM EDTA).

#### Equilibrium Dissociation Constants of IsdB:Hb complexes

The analysis of IsdB:metHb complex series indicates that lowering protein concentration leads to complex dissociation and the formation of free IsdB and Hb dimer molecules. Dissociation constant for the above equilibrium was calculated using a previously reported equation [49], which, in the case of interacting species (called P and R) having the same concentrations, simplifies as follow:

$$H = \frac{\left(2R_{tot} + K_D\right) - \sqrt{K_D\left(4R_{tot} + K_D\right)}}{2R_{tot}} \quad (1)$$

where H is the volume fraction of the higher molecular weight complex, made of an Hb dimer bound by two IsdB molecules (model 2IsdB:$Hb_{dim}$), $R_{tot}$ is the total protein concentration of the sample expressed as concentration of the lower molecular weight complex (here heme concentration), and $K_D$ is the dissociation constant.

Notably, OLIGOMER directly outputs the volume fraction of different components in solution, thus the value corresponding to the 2IsdB:$Hb_{dim}$ has been directly used as the dependent variable.

Analogous approach was applied to IsdB:oxyHb complex to calculate Hb tetramer-dimer equilibrium dissociation constant. In this case, the complex made of two IsdB molecules bound to the $\beta$-chains of an Hb tetramer (2IsdB:$\beta Hb_{tet}$) was considered as H.

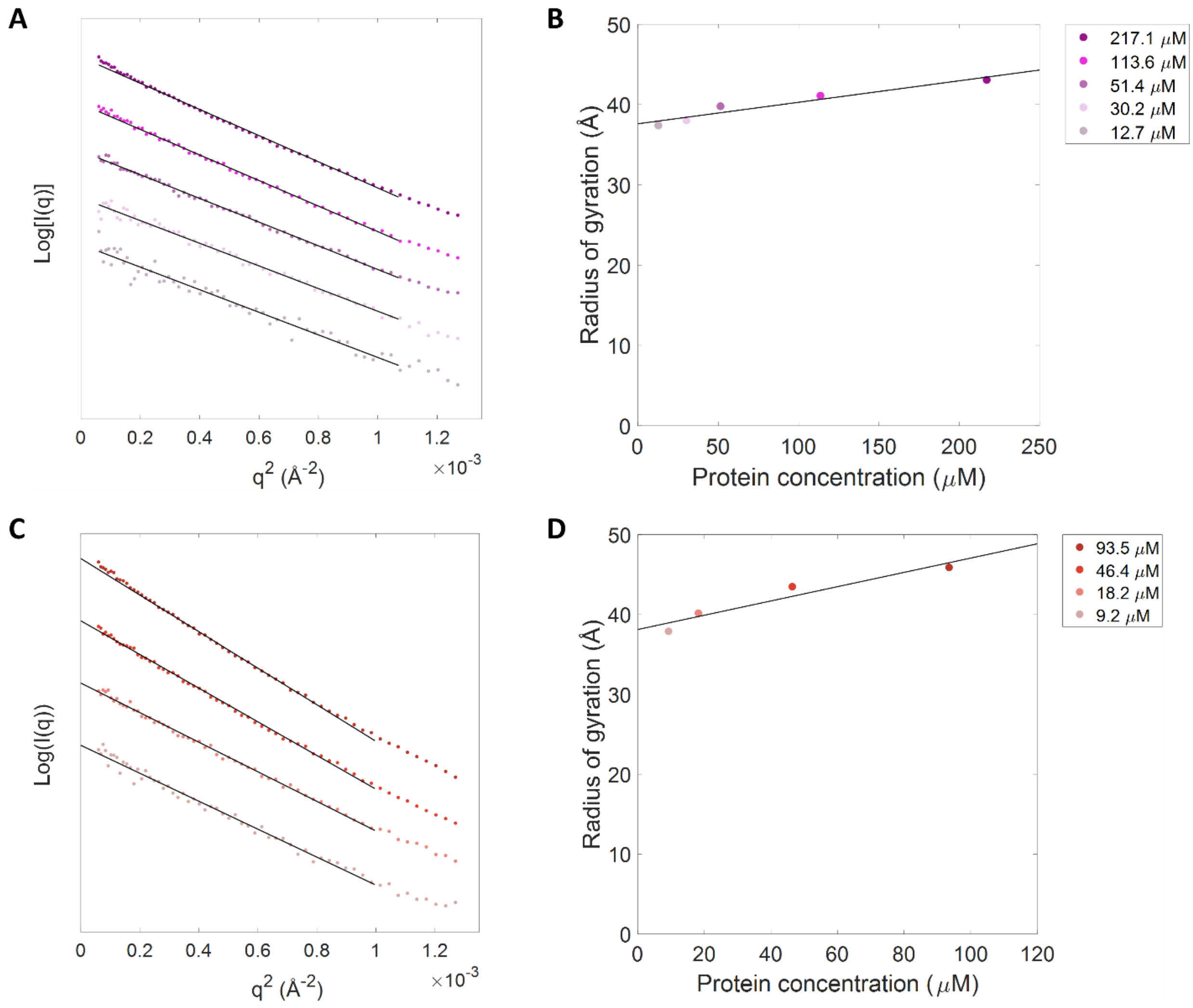


**Fig. S1.** Guinier analysis of IsdB:metHb (panels **A** and **B**) and IsdB:oxyHb (panels **C** and **D**) SAXS data. Black lines in panels a and c represent linear fits in the 0.3/Rg to 1.3/Rg q-range. The radius of gyration extracted from the Guinier analysis is plotted as a function of the complex concentration in panels **B** and **D** (colored dots) together with a linear fit (solid lines).

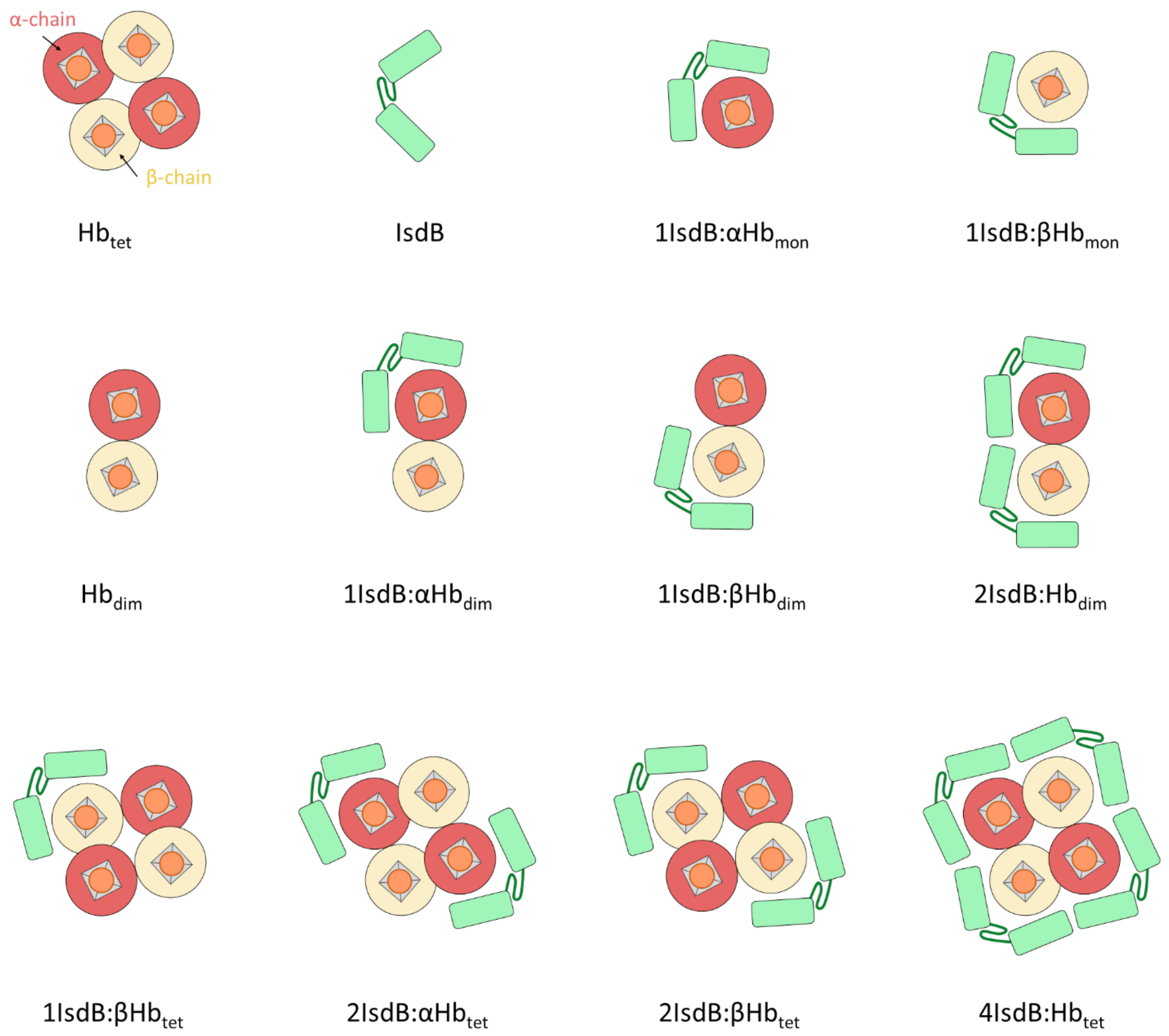


**Fig. S2.** Schematic representation of Hb, IsdB, and all possible *a priori* candidates of the IsdB:Hb complex in solution. As described in the Materials and Methods section, 4IsdB:Hb$_{tet}$ was generated from PDB 5VMM and 3P5Q, and the other models shown here were obtained by removing components from it.

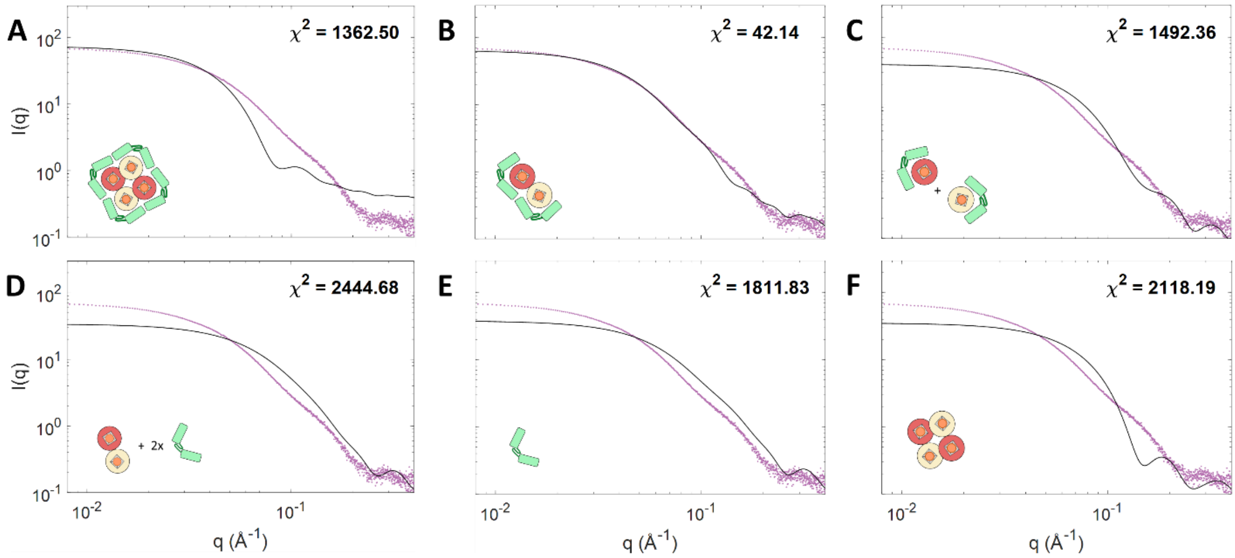


**Fig. S3.** Comparison of I(q) calculated by CRYSOL from the all atom structure (black solid line) and experimental SAXS curves for IsdB:metHb complex at 101.9 $\mu$M (purple dots). SAXS data of IsdB:metHb are compared with theoretical pattern of 4IsdB:$Hb_{tet}$ (**A**), 2IsdB:$Hb_{dim}$ (**B**), linear combination of 1IsdB:α$Hb_{mon}$ + 1IsdB:β$Hb_{mon}$ (**C**), linear combination of $Hb_{dim}$ + 2 IsdB (**D**), IsdB (**E**) and $Hb_{tet}$ (**F**). Theoretical patterns of PDB representing different stoichiometries of the complex were not considered (see Material and Methods).

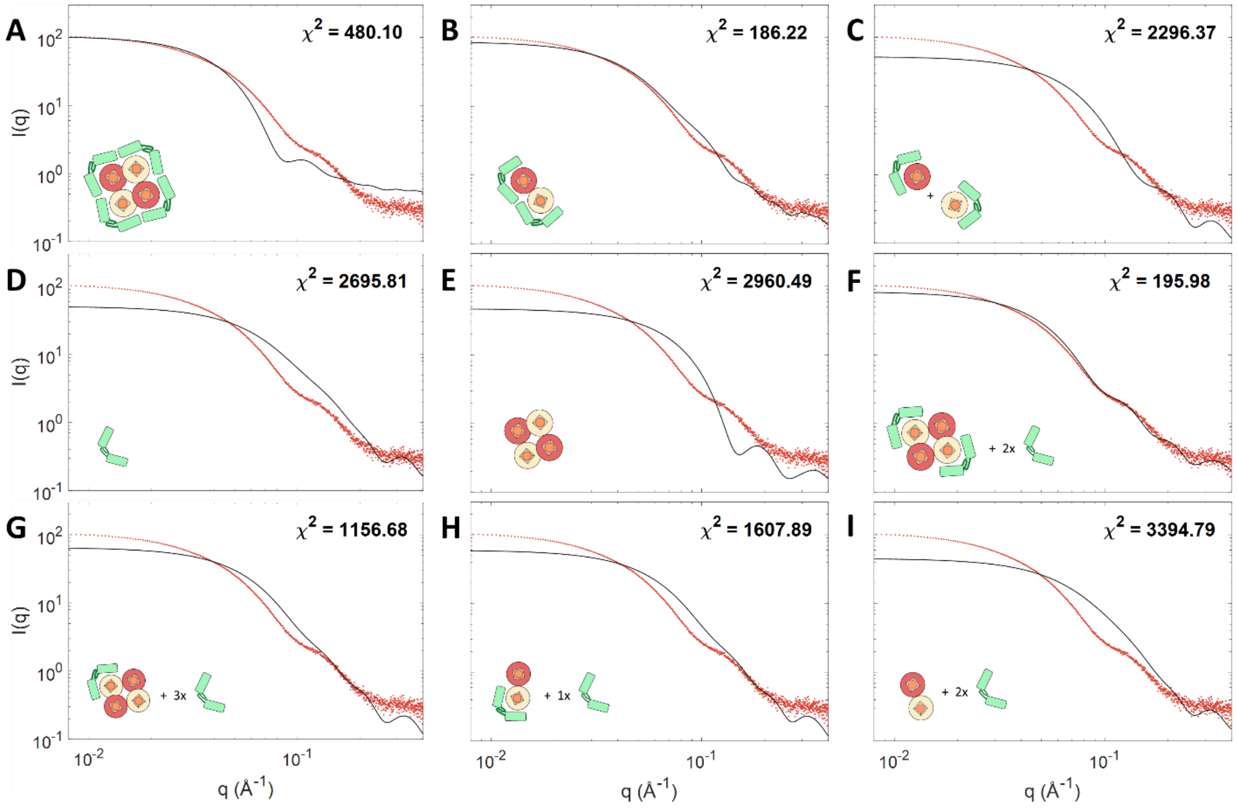


**Fig. S4.** Comparison of I(q) calculated by CRYSOL from the all atom structure (black solid line) and experimental SAXS curves for IsdB:oxyHb complex at 73.5 $\mu$M (red dots). SAXS data of IsdB:oxyHb complex are compared with theoretical pattern of 4IsdB:$Hb_{tet}$ (**A**), 2IsdB:$Hb_{dim}$ (**B**), linear combination of 1IsdB:$\alpha Hb_{mon}$ + 1IsdB:$\beta Hb_{mon}$ (**C**), IsdB (**D**) and $Hb_{tet}$ (**E**), or linear combination of 2IsdB:$\beta Hb_{tet}$ + 2 IsdB (**F**), 1IsdB:$\beta Hb_{tet}$ + 3 IsdB (**G**), 1IsdB:$\beta Hb_{dim}$ + 1 IsdB (**H**), $Hb_{dim}$ + 2 IsdB (**I**). Theoretical patterns of PDB representing 2IsdB:$\alpha Hb_{tet}$ and 1IsdB:$\alpha Hb_{dim}$ were not considered since the preferential binding of IsdB to Hb $\beta$-chains was demonstrated in Fig. S5B.

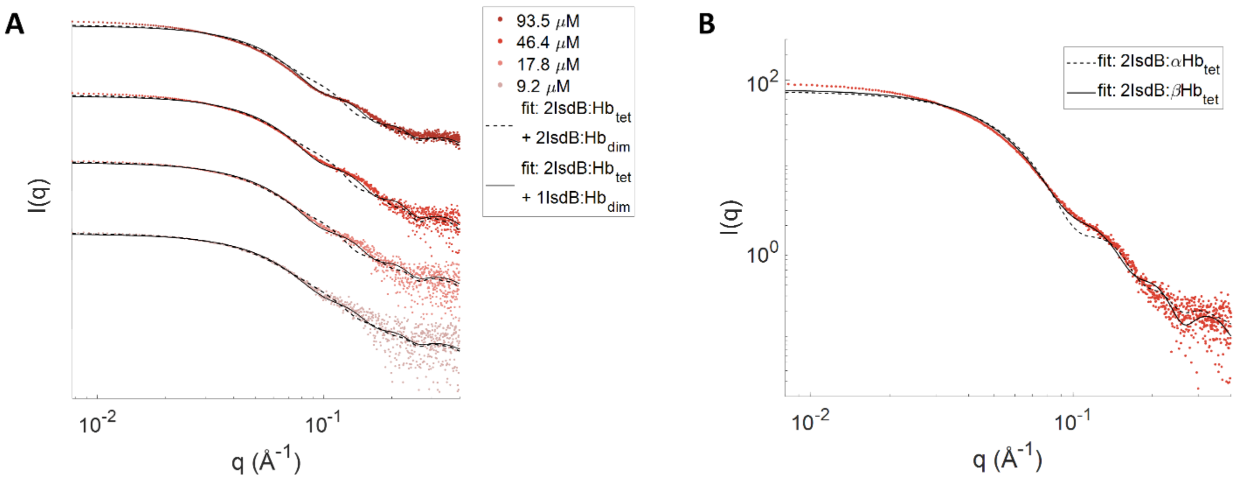


**Fig. S5. A**. SAXS data of the IsdB:oxyHb complex concentration series (dots with the same color code as Fig. 2 and Fig. S1) together with OLIGOMER fitting based on linear combination of 2IsdB:$Hb_{tet}$ and 2IsdB:$Hb_{dim}$ complexes (dashed lines) or 2IsdB:$Hb_{tet}$ and 1IsdB:$Hb_{dim}$ complexes (solid lines). The molar excess of IsdB in solution due to the stoichiometric ratio used to prepare the samples was considered in both OLIGOMER analysis but it has not been inserted in the legend of the figure in order to simplify the comprehension of the plots. **B**. Comparison between SAXS data for IsdB:oxyHb complex at 46.5 $\mu$M (dots) and CRYSOL predictions with IsdB bound to $\alpha$- (dashed, $\chi^2$ = 96.4) or $\beta$- (solid, $\chi^2$ = 49.3) subunits of an Hb tetramer.

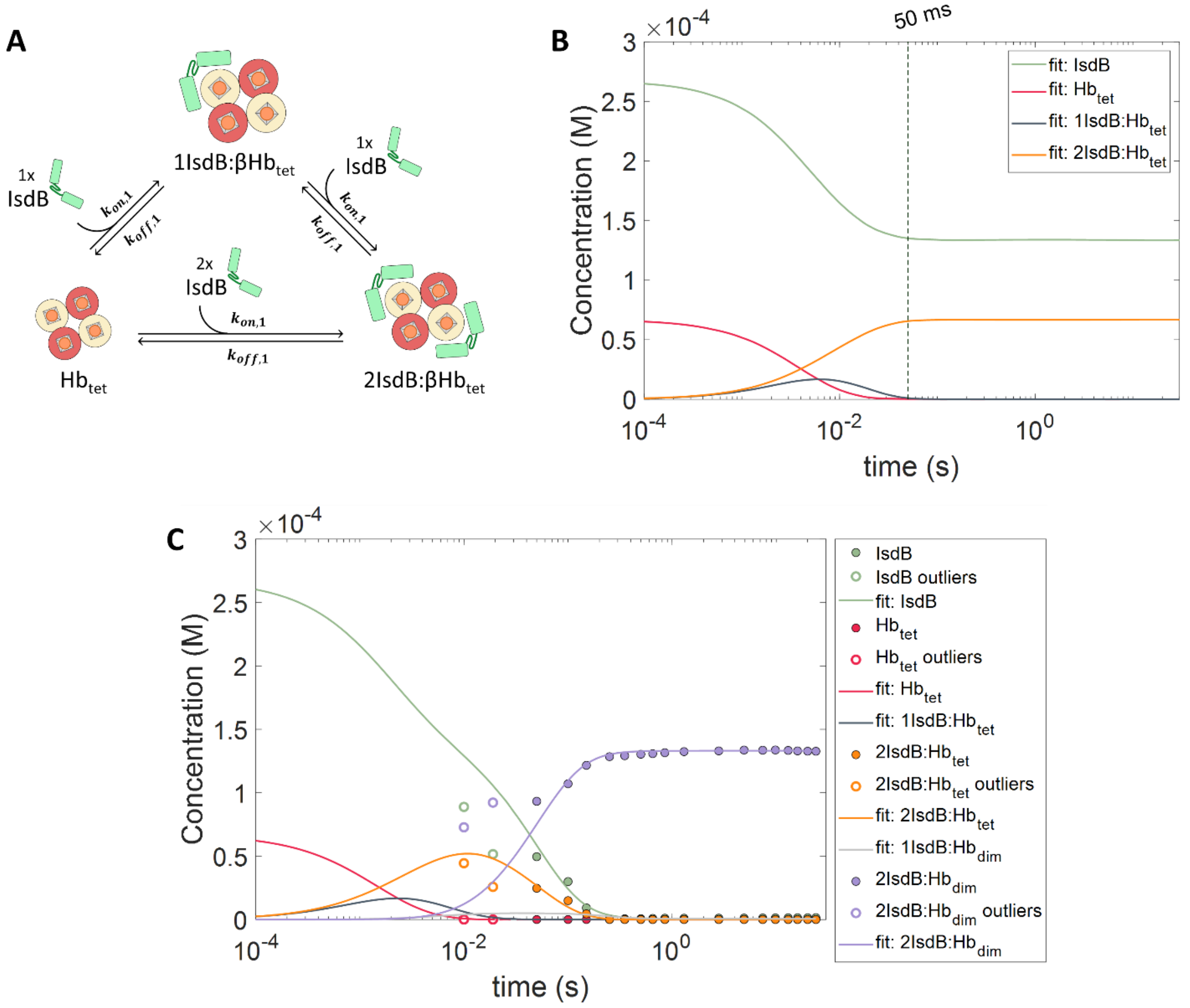


**Fig. S6.** The absence of a static WAXS pattern for IsdB:$\beta$Hb$_{tet}$ complex precludes the chance to correctly analyze the first two time delays of TR-WAXS by linear deconvolution. **A**. Schematic representation of kinetic model used in SimBiology Model Builder to derive differential equations for the molecular processes involved in the IsdB binding to $\beta$-chains of an Hb tetramer. **B**. Simulated concentration of intermediate species in 2IsdB:$\beta$Hb$_{tet}$ complex formation as a function of time. Data were calculated using the kinetic model defined in (**A**) and published SPR kinetic constants ($k_{on} = 5.30 \cdot 10^5$ $M^{-1}$ $s^{-1}$ and $k_{off} = 3.7 \cdot 10^{-2}$ $s^{-1}$, [7]) in SimBiology Model Analyzer. Color scheme is consistent with Fig. 5. The plot shows that only after 50 ms (vertical dashed line) the 1IsdB:$\beta$Hb$_{tet}$ complex concentration becomes negligible. **C**. Representation of species in IsdB:Hb complex formation estimated from the linear combination of TR-WAXS data for time delays outliers (*i.e* 10 ms and 18 ms, open circles) in comparison with species (closed circles) and SimBiology fitting (solid lines) as reported in Fig. 5.

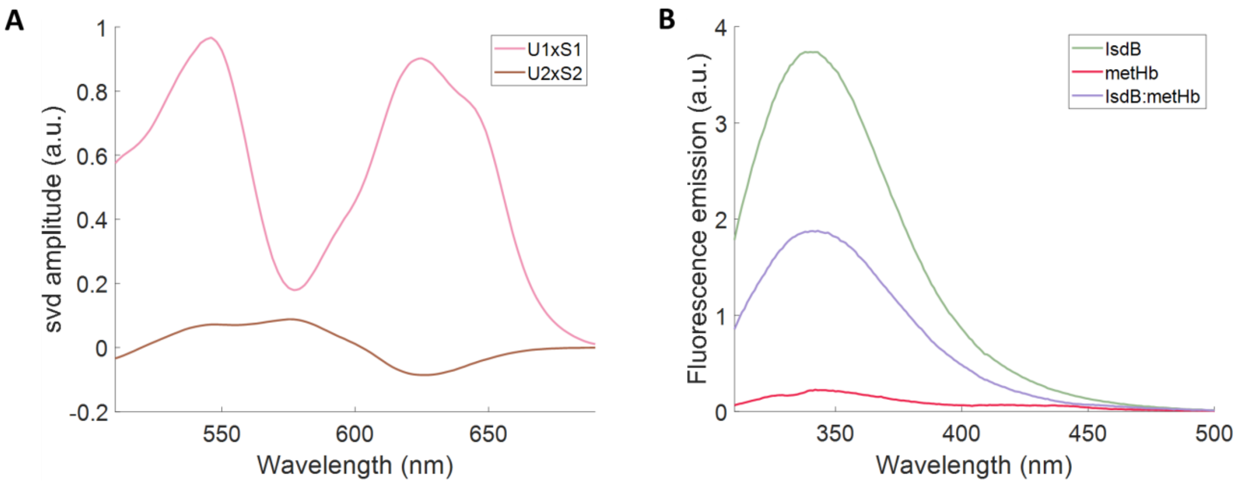


**Fig. S7. A**. Main components from the singular value decomposition of TR-OA. Color scheme is consistent with Fig. 6B. Interaction between IsdB and metHb was evaluated by exploiting the fluorescence of three IsdB tryptophan residues (W195 on NEAT1 domain, W392 on NEAT2 domain and an additional tryptophan on the recombinantly-added StrepTag®II). Indeed, the fluorescence emission of Hb tryptophans is quenched by the heme cofactor. Emission spectra of isolated IsdB (green line) and metHb (red line) or their mixture (purple line) in a stoichiometric amount at 5 µM concentration were recorded from 310 nm to 500 nm at 20 °C in the experimental buffer with excitation at 298 nm using a HORIBA FluoroMax-3 spectrofluorometer.

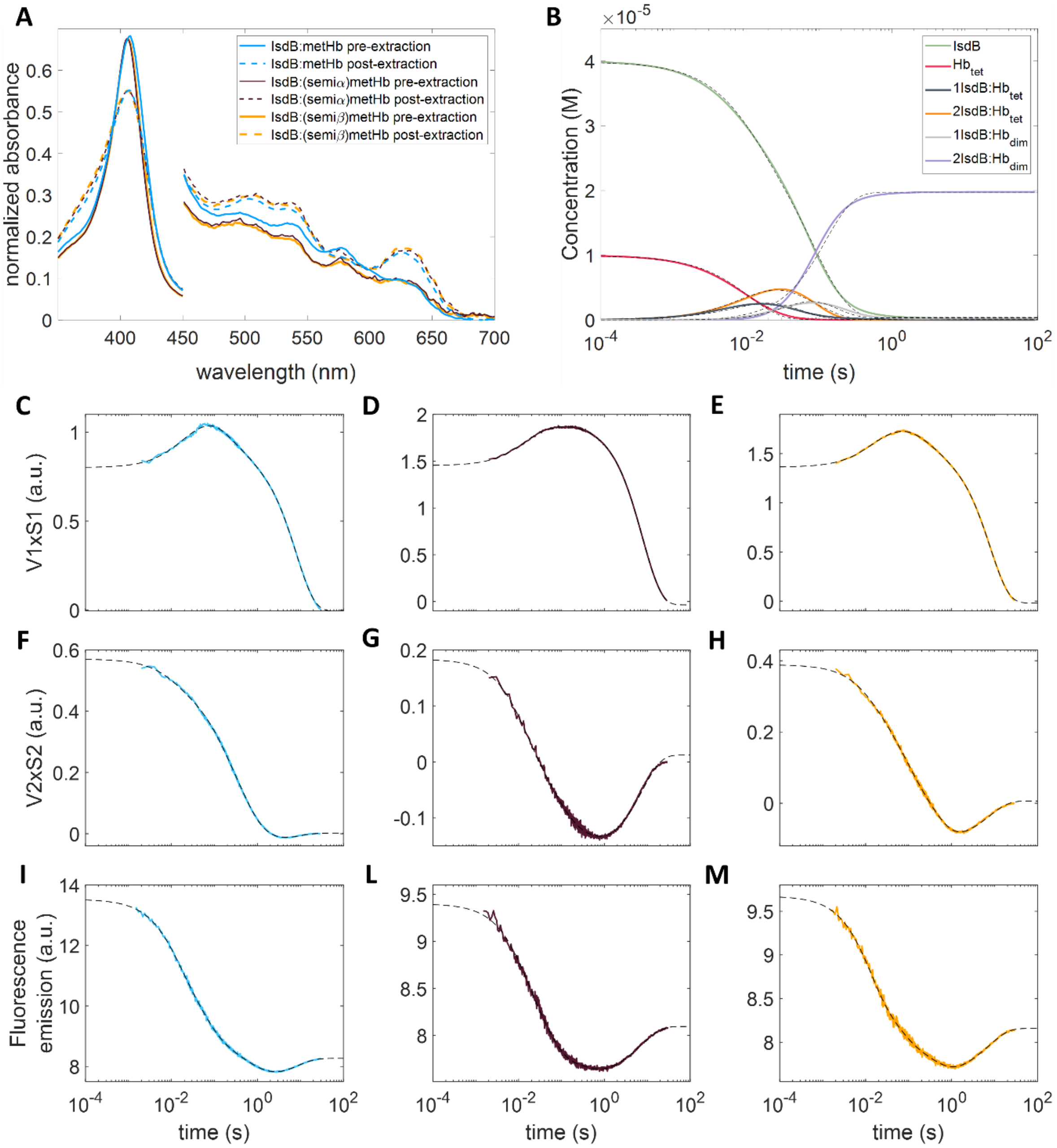


**Fig. S8. A**. Absorption spectra of IsdB mixed with metHb or semi-metHbs. Pre-extraction spectra (solid lines) are normalized on the basis of maximum absorbance of sample with native metHb, and post-extraction spectra (dashed lines) are normalized accordingly. The good overlap among post-extraction spectra suggests that the heme transfer process occurs in a comparable manner either when globins are fully (e.g. native metHb) or half-saturated (e.g. semi-metHbs) with the cofactor. Absorption values from 450 to 700 nm are multiplied by a factor 5 to improve visualization of Hb Q bands. **B-M**. Global fitting (black dashed lines) as a multiexponential function of the time course of simulated SimBiology traces for isolated proteins, intermediate, and final complexes at a concentration of IsdB and metHb chains of 40 µM (**B**). The previously defined kinetic model in SimBiology and the extrapolated microscopic constants (**Table 1**) were exploited to simulate this time course of molecular species within IsdB:Hb interaction at the concentration used for the experiments with native and semiHbs. Spectroscopic signals probing the interaction of

IsdB with native (cyan), semi($\alpha$) (purple), and semi($\beta$) (orange) forms of metHb at the same concentration (**C-M**). Signals include first (**C-E**) and second (**F-H**) SVD components from absorption spectroscopy and fluorescence signals (**I-M**), which were already used to study the IsdB:metHb interaction at the concentration comparable to TR-WAXS experiments. Fitting was performed by adapting the lsqmultinonlin function [51] in MATLAB. The good quality of the fitting confirms that molecular events probed by spectroscopy are well described by the SimBiology simulation, supporting the reliability of the defined model and the extrapolated microscopic constants. Moreover, the analysis quantitatively demonstrates that metHb and semiHbs share the same kinetic behavior, although spectroscopic signals and hence amplitudes can differ due to the effect of the lacking heme chromophore. The results obtained with semiHbs are in agreement with previous SPR experiments, when taking into account that IsdB always first binds to beta subunits of Hb, even though in the apo form [7].

Data were well represented by 6 exponentials with respect to 7 in the TR-WAXS conditions dataset. While r5 and r6 are comparable with r6 and r7 in previous analysis, as they are expected to be not concentration-dependent, all these kinetics lacked a time constant attributed to dimerization/activation (r4 in TR-WAXS conditions dataset). This can be explained by the fact that a lower concentration of reactant makes bimolecular reaction slower (see r1, r2 and r3) and this process is no longer a rate limiting step in binding to α-chains (**Table S2**).

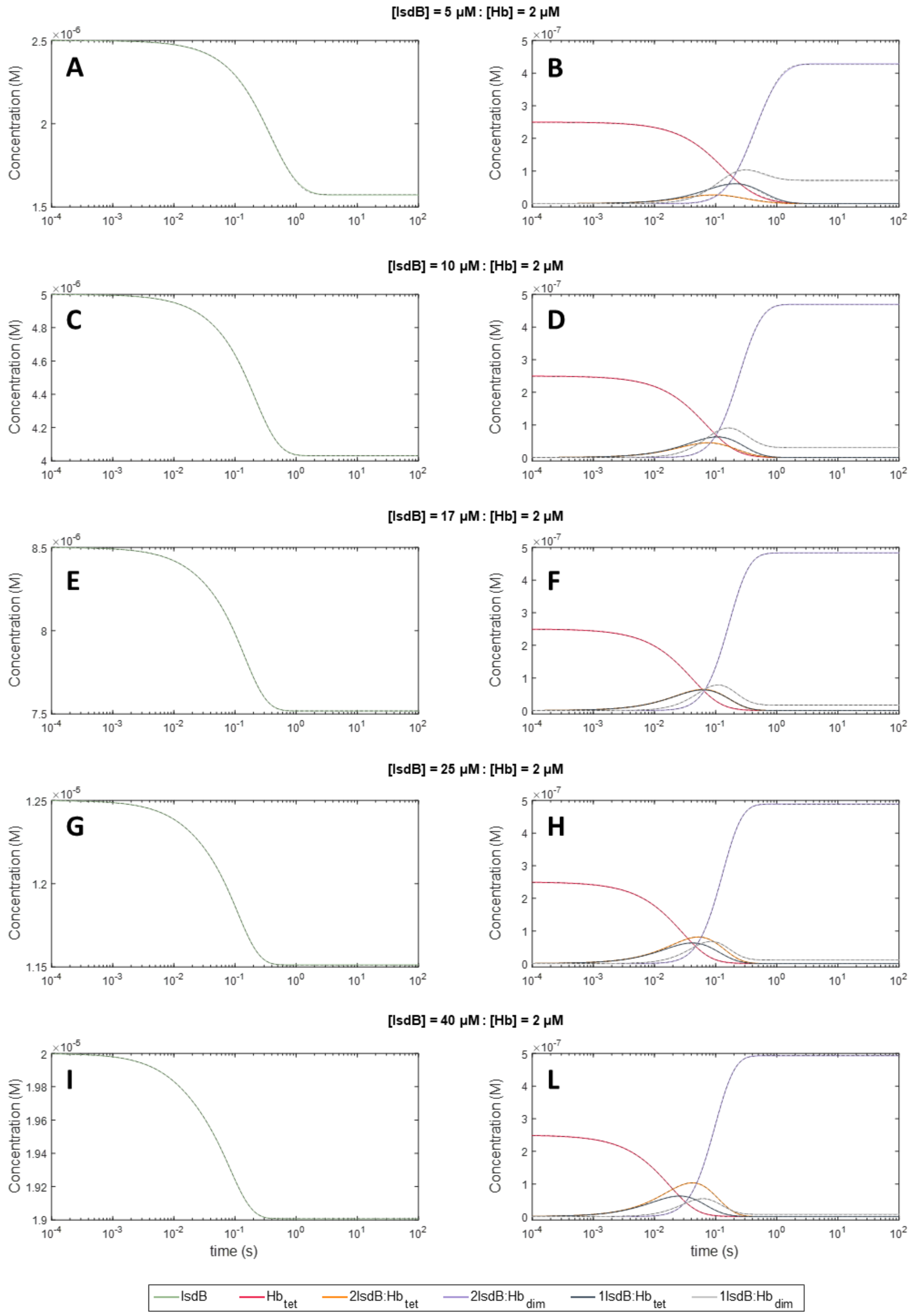


**Fig. S9.** Simulation of concentration profiles of molecular species (color scheme is kept comparable with respect to Fig. 6) within IsdB:Hb PPI based on the kinetic model in Fig. 5C, rate constants in Table 1, and the experimental conditions of a published experiment by Bowden and co-workers [6]. In the reference paper, the authors studied the IsdB:Hb PPI following the spectroscopic signal of heme transfer at 406 nm by rapidly mixing 2 μM metHb (heme concentration) and increasing concentration of IsdB (from 2.5 to 40 µM), and fitting the signal as a sum of four exponentials (observed kinetic constants are reported in the table at the bottom of the Table S3). The concentration profiles were globally fitted as a sum of three exponential functions according to the analysis reported in Fig. 6 and S6. As already observed in Table S1 and S2, the fastest molecular process (*i.e.* r1) is not detectable by absorption spectroscopy, and aside from that, rate constants from SimBiology simulations and published observed kinetic rates appear in good agreement confirming the reliability of the kinetic model defined in Fig. 5C and rate constants in Table 1 in representing the evolution of molecular species over the time of the IsdB:Hb PPI.

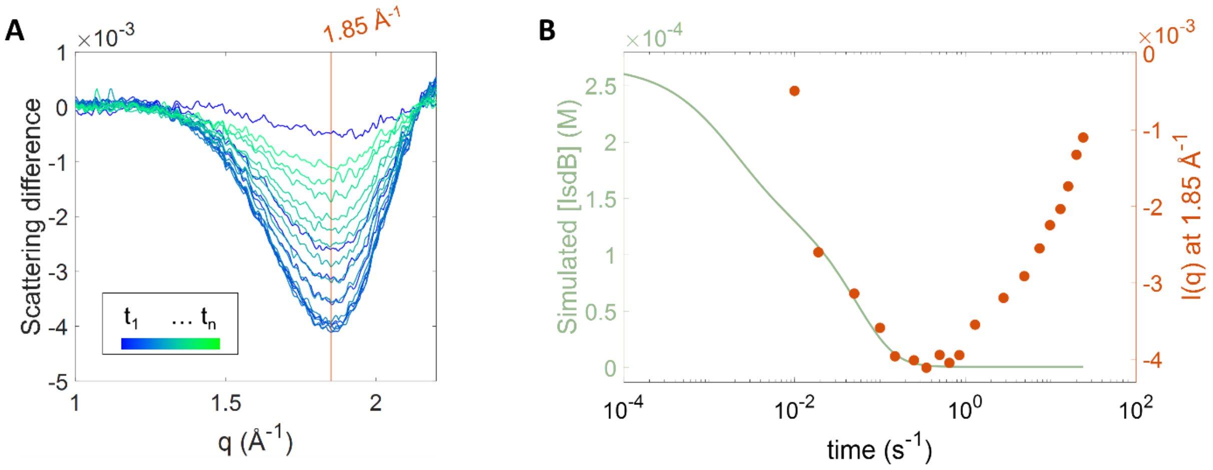


**Fig. S10.** Thermal effect of the IsdB:Hb complex formation on solvent scattering. **A**. TR-WAXS difference patterns around the water scattering peak (2 $Å^{-1}$) collected at different time-delays from mixing. **B**. Time-evolution of the signal amplitude at q = 1.85 $Å^{-1}$ (red circles) and of the IsdB concentration obtained from the simulation described in the main text (green line). The water scattering peak is sensitive to the local temperature in the volume probed by the X-ray beam [13, 27]. Accordingly, data indicate that an initial fast heating phase is followed by a slower sample cooling. The initial heating phase suggests that heat is released to the solvent upon binding of IsdB to metHb in agreement with recent isothermal thermal calorimetry measurements [6, 38]. Indeed, if properly scaled, the fast phase of the signal probing the solvent temperature can be superimposed to the time decay of the IsdB concentration in solution, which is due to the IsdB:metHb complex formation. The following cooling phase, occurring between a few hundred milliseconds and a few seconds, can be ascribed to heat diffusion out of the volume probed by the X-ray beam.

**Table S1.** Apparent shared rate constants (r) from the fitting as a sum of exponential functions of SimBiology simulated and absorbance datasets of the reaction between IsdB and metHb at 250 µM (a concentration similar to TR-WAXS experiment). Cells are colored based on normalized amplitudes of exponential decay (blue) or rise up (red). Graphical representation of the fit is shown in Fig. 6. * The number of traces describing the earlier phase is small and does not allow the accurate fit convergence.

| | **r1 ($s^{-1}$)** | **r2 ($s^{-1}$)** | **r3 ($s^{-1}$)** | **r4 ($s^{-1}$)** | **r5 ($s^{-1}$)** | **r6 ($s^{-1}$)** | **r7 ($s^{-1}$)** |
|---|---|---|---|---|---|---|---|
| | 811 ± 223 x$10^3$ * | 374 ± 64 | 98.6 ± 2.8 | 15.0 ± 0.5 | 2.59 ± 0.32 | 0.84 ± 0.11 | 0.113 ± 0.004 |
| | **Amplitude %** | **Amplitude %** | **Amplitude %** | **Amplitude %** | **Amplitude %** | **Amplitude %** | **Amplitude %** |
| SimBiology IsdB | 23 | 10 | 18 | 49 | -1 | 0 | 0 |
| SimBiology $Hb_{tet}$ | 62 | 33 | 5 | -1 | 0 | 0 | 0 |
| SimBiology 1IsdB:$Hb_{tet}$ | -96 | 47 | 52 | -4 | 1 | 0 | 0 |
| SimBiology 2IsdB:$Hb_{tet}$ | 0 | -79 | -14 | 100 | -7 | 0 | 0 |
| SimBiology 1IsdB:$Hb_{dim}$ | 0 | 6 | -100 | 31 | 55 | 0 | 0 |
| SimBiology 2IsdB:$Hb_{dim}$ | 0 | 9 | -8 | -92 | 4 | 0 | 0 |
| SVD V1xS1 IsdB:metHb | 0 | 5 | -1 | 9 | 9 | 13 | 65 |
| SVD V2xS2 IsdB:metHb | 0 | 23 | 10 | 34 | 25 | 8 | -23 |
| Fluo ex 298 nm IsdB:metHb | 0 | 27 | 42 | 16 | 6 | 5 | 4 |

**Table S2.** Apparent shared rate constants from the fitting as a sum of exponential functions of SimBiology simulated and absorbance datasets of the reaction between IsdB and metHb at 40 µM, together with absorbance datasets of the reaction between IsdB and (semi)metHb at the same concentration. Cells are colored based on normalized amplitudes of exponential decay (blue) or rise up (red). Graphical representation of the fit is shown in Fig. S6.

| | | r1 ($s^{-1}$) | r2 ($s^{-1}$) | r3 ($s^{-1}$) | r4 ($s^{-1}$) | r5 ($s^{-1}$) | r6 ($s^{-1}$) |
|---|---|---|---|---|---|---|---|
| | | 92.8 ± 2.1 | 23.1 ± 1.3 | 10.1 ± 1.3 | 2.39 ± 0.16 | 0.80 ± 0.04 | 0.128 ± 0.008 |
| | | **Amplitude %** | **Amplitude %** | **Amplitude %** | **Amplitude %** | **Amplitude %** | **Amplitude %** |
| Simbiology | IsdB | 28 | 10 | 62 | 0 | 0 | 0 |
| | $Hb_{tet}$ | 100 | 0 | 0 | 0 | 0 | 0 |
| | 1IsdB:$Hb_{tet}$ | -99 | 100 | 0 | 0 | 0 | 0 |
| | 2IsdB:$Hb_{tet}$ | -100 | 19 | 81 | 0 | 0 | 0 |
| | 1IsdB:$Hb_{dim}$ | 14 | -100 | 84 | 0 | 0 | 0 |
| | 2IsdB:$Hb_{dim}$ | 0 | 27 | -100 | 0 | 0 | 0 |
| SVD V1xS1 | IsdB:metHb | -8 | -26 | 15 | 12 | 0 | 73 |
| | IsdB:(semiα)metHb | -15 | -8 | 0 | 0 | 0 | 100 |
| | IsdB:(semiβ)metHb | -12 | -16 | 8 | 11 | 0 | 81 |
| SVD V2xS2 | IsdB:metHb | 14 | 0 | 23 | 47 | 16 | -5 |
| | IsdB:(semiα)metHb | 38 | 22 | 25 | 16 | 0 | -50 |
| | IsdB:(semiβ)metHb | 16 | 20 | 22 | 42 | 0 | -23 |
| Fluo ex 298 nm | IsdB:metHb | 32 | 31 | 14 | 12 | 11 | -12 |
| | IsdB:(semiα)metHb | 37 | 48 | 2 | 13 | -4 | -27 |
| | IsdB:(semiβ)metHb | 49 | 23 | 8 | 20 | -3 | -24 |

**Table S3.** Comparison of observed kinetic constants from Bowden and co-workers [6] and the rate constants calculated on the basis of SimBiology simulation. * The r6 and r7 constants were picked from the global fitting in Fig. 6 as they are assumed to be concentration independent.

| | **[IsdB] = 5 µM** | **[IsdB] = 10 µM** | **[IsdB] = 17 µM** | **[IsdB] = 25 µM** | **[IsdB] = 40 µM** |
|---|---|---|---|---|---|
| Global fit: r1 ($s^{-1}$) | 26.46 ± 0.88 | 30.64 ± 0.34 | 33.23 ± 0.14 | 40.61 ± 0.09 | 59.99 ± 0.14 |
| Global fit: r2 ($s^{-1}$) | 9.71 ± 0.12 | 16.14 ± 0.10 | 24.09 ± 0.08 | 29.09 ± 0.06 | 35.86 ± 0.10 |
| *Bowden: $k_{1,obs}$ ($s^{-1}$)* | *11 ± 2* | *25 ± 2* | *32 ± 3* | *49 ± 7* | *108 ± 8* |
| Global fit: r3 ($s^{-1}$) | 2.41 ± 0.01 | 5.35 ± 0.01 | 9.11 ± 0.01 | 12.59 ± 0.01 | 16.48 ± 0.01 |
| *Bowden: $k_{2,obs}$ ($s^{-1}$)* | *1.9 ± 0.2* | *2.1 ± 0.2* | *5 ± 1* | *4 ± 1* | *4 ± 1* |
| Global fit: r6* ($s^{-1}$) | 0.84* ± 0.11 | 0.84* ± 0.11 | 0.84* ± 0.11 | 0.84* ± 0.11 | 0.84* ± 0.11 |
| *Bowden: $k_{3,obs}$ ($s^{-1}$)* | *0.41 ± 0.03* | *0.42 ± 0.05* | *0.5 ± 0.1* | *0.5 ± 0.1* | *0.5 ± 0.1* |
| Global fit: r7* ($s^{-1}$) | 0.113* ± 0.004 | 0.113* ± 0.004 | 0.113* ± 0.004 | 0.113* ± 0.004 | 0.113* ± 0.004 |
| *Bowden: $k_{4,obs}$ ($s^{-1}$)* | *0.12 ± 0.02* | *0.13 ± 0.03* | *0.14 ± 0.04* | *0.15 ± 0.03* | *0.13 ± 0.04* |
